# Material Composition and Implantation Site Affect in vivo Device Degradation Rate

**DOI:** 10.1101/2024.09.09.612079

**Authors:** K. M. Pawelec, J. M.L. Hix, A. Troia, M. Kiupel, E. M. Shapiro

## Abstract

Successful tissue engineering requires biomedical devices that initially stabilize wounds, then degrade as tissue is regenerated. However, the material degradation rates reported in literature are often conflicting. Incorporation of in situ monitoring functionality into implanted devices would allow real time assessment of degradation and potential failure. This necessitates introduction of contrast agent as most biomedical devices are composed of polymeric materials with no inherent contrast in medical imaging modalities. In the present study, computed tomography (CT)-visible radiopaque composites were created by adding 5-20wt% tantalum oxide (TaO_x_) nanoparticles into polymers with distinct degradation profiles: polycaprolactone (PCL), poly(lactide-co-glycolide) (PLGA) 85:15 and PLGA 50:50, representing slow, medium and fast degrading materials respectively. Radiopaque phantoms, mimicking porous tissue engineering devices, were implanted into mice intramuscularly or intraperitoneally, and monitored via CT over 20 weeks. Changes in phantom volume, including collapse and swelling, were visualized over time. Phantom degradation profile was determined by polymer matrix, regardless of nanoparticle addition and foreign body response was dictated by the implant site. In addition, degradation kinetics were significantly affected in mid-degrading materials, transitioning from linear degradation intramuscularly to exponential degradation intraperitoneally, due to differences in inflammatory responses and fluid flow. Nanoparticle excretion from degraded phantoms lagged behind polymer, and future studies will modulate nanoparticle clearance. Utilizing in situ monitoring, this study seeks to unify literature and facilitate better tissue engineering devices, by highlighting the relative effect of composition and implant site on important materials properties.

## 1 Introduction

As the field of tissue engineering has grown, so has the number of available materials for use and the complexity of potential implanted devices to stimulate the repair of damaged and diseased tissues. Despite these advances, the translation of novel materials and devices to the clinic remains challenging. This is in part due to a lack of understanding around how materials interact dynamically within the physiological environment, which feature constantly changing flows of interstitial fluids, soluble molecules, proteases and cellular components. In this environment, the precise structural properties reported in vitro broaden, making prediction of functional efficacy difficult. Added to design uncertainty, the clinical use of devices introduces risks to patients in the form of unanticipated movement of the device, fracture, and infection [1]. All of this is prompting the trend towards incorporating functionality for in situ monitoring into implantable devices.

Biomedical imaging techniques are an emerging way to gain information on both biological responses to devices and changes experienced by the implanted devices [2]. In preclinical stages of design, following biological responses over time can reduce animal number while creating an integrated picture of the stages of tissue repair, from acute post-implant inflammation, to extra cellular matrix (ECM) deposition, vascularization, and mature functional tissue formation. However, to fully integrate our understanding of biological responses around tissue engineering devices, changes to the device itself, such as collapse, damage and degradation, should be tracked alongside these signals. Degradation, in particular, has proven to be a key property that is difficult to determine for many materials, with in vitro measurements generally underestimating in vivo degradation rates, due to a lack of fluid flow and inflammatory factors [3–4].

While the monitoring of tissue engineering devices could add considerably to the understanding of how these devices function, it represents a significant challenge as the majority of implanted devices are made from polymers, which have no inherent contrast from native tissue post-implantation. To characterize polymer devices after implantation, some form of imaging contrast agent must be added into the device [5]. Ideally, this contrast agent would fulfill a number of requirements, including: low systemic toxicity, no interference with tissue regeneration, remaining associated with the device structure for its lifetime of use, and not introducing unwanted artifacts into other imaging modalities to allow for multimodal imaging of distinct biological processes.

Identifying a potential contrast agent for incorporation into polymers first requires a selection of imaging modality. Available techniques include fluorescence, positron emission tomography (PET), magnetic resonance imaging (MRI) and computed tomography (CT) [2]. These techniques vary in resolution, their ability to provide three-dimensional anatomical data, and the depth of the tissue that can be scanned. Indeed, a variety of imaging modalities have been utilized to track polymer degradation post-implantation, including fluorescence [6], MRI [7] and CT [8–9]. Contrast agents can also be combined for imaging the same material using multiple modalities [10]. While MRI has the greatest resolution for 3D imaging of soft tissues, CT technologies remain the workhorse of clinical imaging for their ease of use, relative cost-effectiveness and good resolution. In addition, it is easy to translate CT imaging across the spectrum from the lab to preclinical studies and finally clinical practice, and for these reasons, this study focuses on radiopaque contrast agent addition [11].

Introduction of x-ray attenuating contrast into model implantable biomedical devices (phantoms) has been accomplished in a variety of ways. The polymer chain can be chemically modified through the introduction of radiopaque elements, such as iodine groups [12–13]. However, there are tradeoffs to modification of the polymer backbone, as it can also affect the degradation profile of the material and degradation by-products [14]. Addition of radiopaque nanoparticles offer another way to incorporate contrast that allows for independent tuning of the polymer matrix and the nanoparticle agent, and could be further modified as drug delivery vehicles [15]. Radiopaque nanoparticles include noble metals like gold [16], and metal oxides, including bismuth [17] and tantalum [18]. Metal oxides have a clinical advantage in their cost-effectiveness and in the case of tantalum oxide, have demonstrated biocompatibility and superior attenuation over iodinated compounds [19–20].

Contrast agents have been used to follow in vitro degradation profiles of devices over time, highlighting differences in degradation kinetics leading to structural device collapse [10,21]. In vivo, the majority of studies following degradation have been performed on hydrogel systems, which have relatively quick degradation kinetics and do not represent the spectrum of clinical devices currently in use. This study extends these findings by examining the degradation of synthetic polymers with degradation kinetics ranging from weeks to years. The addition of tantalum oxide (TaO_x_) nanoparticles was used to create radiopaque phantom devices and the effect of the composite composition, including both polymer matrix and amount of nanoparticles present, was studied alongside the effects of implantation site (intramuscular or intraperitoneal). This study shows successful in vivo tracking of phantom devices out to 20 weeks post-implantation utilizing micro-CT (μCT). Further, it demonstrates that both composition and implant site affect device degradation, highlighting that capability for in situ monitoring will be a critical component in the clinical translation of future tissue engineering devices.

## 2 Results

The ability to monitor biomedical implants is necessary for the early detection of defects, tears and movement that can lead to catastrophic device failure in patients and at a minimum delay treatment, prolonging patient suffering [5]. In the field of tissue engineering, devices are designed to degrade over time, allowing newly regenerated tissue to replace a supporting structure gradually. Achieving this requires a match of mechanical properties and degradation profile to tissue regeneration, to promote tissue in-growth [22]. However, degradation profiles in vitro do not match in vivo rates and to-date the preferred method of measuring the degradation of implanted biomedical devices is via destructive testing at discrete time points [4]. For multifactor studies this becomes unwieldy very quickly, necessitating transition to in situ monitoring. However, as the porosity and shape of implanted devices can affect their degradation profile, it imperative that the incorporation of contrast agents does not affect the morphology of the devices [23].

Radiopaque composites were created by homogeneously incorporating a hydrophobic TaO_x_ nanoparticle into hydrophobic polymer matrices [21]. To mimic tissue engineering devices, porous phantoms were created with micro-scale porosity (< 100μm) to accommodate nutrient diffusion and macro-scale porosity (200 - 500 μm) for cell and tissue infiltration. Utilizing a salt leaching method, devices were produced from three polymer matrices: polycaprolactone (PCL), poly(lactide-co-glycolide) (PLGA) 85:15 and PLGA 50:50. While all three are hydrophobic polyesters, they vary in their degradation rate. The fastest degrading polymer, PLGA 50:50, hydrolyzes in weeks, while PLGA 85:15 requires months to degrade [4]. In contrast, PCL degrades over 2-3 years, making it essentially non-degradable for the duration of this 20 week study [24]. By using different polymer matrices, we demonstrate the ability to vary the degradation rate of phantoms and X-ray attenuation independently.

Each polymer matrix, incorporating up to 20wt% TaO_x_ nanoparticles, has been shown previously to produce phantoms with identical porous architectures, Figure 1(a-d), and no effect of nanoparticle addition on mechanical properties [21,25]. The nominal incorporation of TaO_x_ addition was kept between 5-20wt%, as this is a range shown to have X-ray attenuation above background tissue, and be biocompatible with a variety of primary cell types, including macrophages and glial cells [26–27]. All composites incorporating nanoparticles had homogeneous X-ray attenuation that scaled with TaO_x_ addition, from 350.5 ± 63.1 HU to 803 ± 131 HU for device matrices incorporating 5 and 20wt% TaO_x_ respectively. This correlated to 5.98 ± 0.83 wt% and 16.84 ± 0.56 wt% TaO_x_ nanoparticles when measured via thermogravimetric analysis (Supplemental Table S1). Tuning radiopacity via nanoparticle incorporation has been utilized in a variety of systems, including bismuth oxide nanoparticles [17] and gold nanoparticles [8]. In the present case, the measured radiopacity per weight percent TaO_x_ addition is lower than other studies, as the measured attenuation is a composite of the radiopacity of the phantom walls and the aqueous buffer perfusing the highly porous matrix.

**Figure 1:**
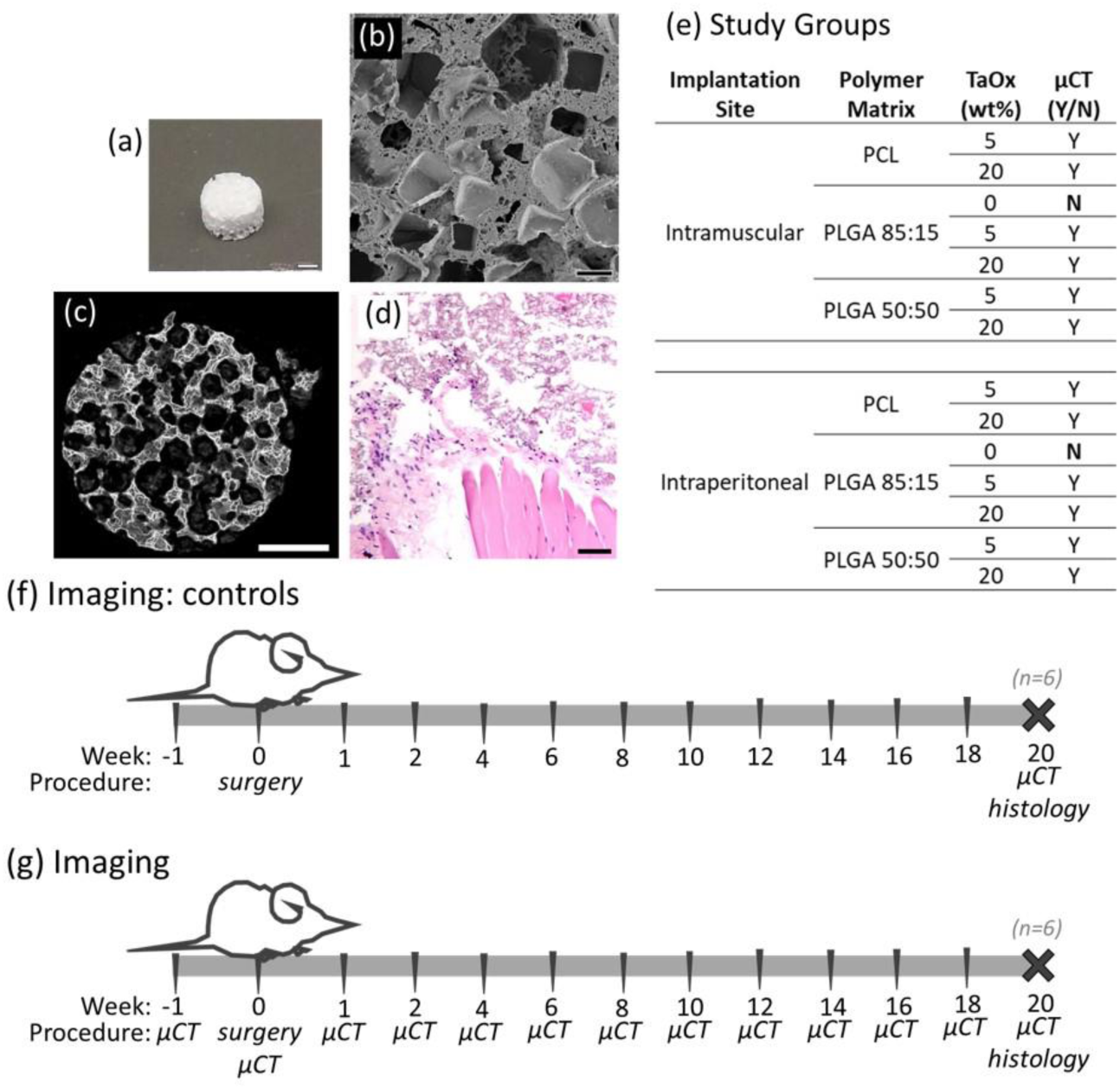
Radiopaque phantoms were created from composites of biocompatible polymers and TaO_x_ nanoparticles for study of in vivo degradation. Implanted phantoms mimicked porous tissue engineering devices as seen (a) macroscopically, (b) in scanning electron microscopy and (c) a 3D rendering of μCT imaging. (d) The open porous structure was still visible after phantoms were placed in vivo, as shown on day 1 post- implantation in the intraperitoneal site with H&E staining. (e) Degradation of radiopaque phantoms was tracked as a function of phantom composition and implant site. A schematic of the 20 week study are shown in (f) for controls without in situ monitoring and (g) all other groups. Scale bar (a, c): 1 mm, (b) 200 μm, (d) 15 μm.

### 2.1 Serial Imaging Longer Term

In vitro, radiopaque composites have degradation rates dictated by the polymeric matrix, which can range from weeks (PLGA 50:50), to months (PLGA 85:15) to years (PCL) [3,21,24]. While in vivo degradation rates are known to differ from in vitro, it is not known to what extent this variation is due to fluid flow, mechanical deformation or inflammatory responses, all of which are affected by the physiological environment. Therefore, to assess the universality of device degradation kinetics, radiopaque phantoms were implanted into mice intramuscularly and intraperitoneally and serially monitored over 20 weeks via μCT, Figure 1(e-g). As a control for degradation kinetics and any alteration in the foreign body response with respect to TaO_x_ nanoparticles, phantoms without nanoparticle addition (0wt% TaO_x_) or serial imaging were used. Together, this allowed for comprehensive picture of how phantom material composition and implantation site might affect tissue regeneration strategies.

Regardless of implantation site, intramuscular or intraperitoneal, serial monitoring of implanted phantoms was successfully performed over 20 weeks post-implantation. At no point in the study was there was an exceptionally fast release of TaO_x_ nanoparticles from the phantoms that obscured phantom features during μCT imaging or made it difficult to obtain metrics describing degradation. This is despite significantly different rates of phantom degradation exhibited by the various polymer composites. Overall, phantoms implanted intramuscularly incorporating 20wt% TaO_x_ had features with the greatest perceived resolution. This was in part due to the ability to restrict movement of the implant site (hind leg) compared to the intraperitoneal site, which is affected by the involuntary movement of breathing. With the addition of 5wt% TaO_x_, only gross scaffold features could be quantified, such as total phantom volume, but separating out the matrix from tissue proved challenging for automated imaging. Therefore, direct information on the matrix was limited to phantoms incorporating 20wt% TaO_x_ nanoparticles. This follows what has been shown previously, where a minimum concentration of contrast agent exists that facilitates measurement of fine features in devices, due to background noise dominating device segmentation from surrounding tissue [11].

Each type of polymer matrix had a distinct degradation response that was clearly discernible via serial monitoring. This was illustrated by matrices incorporating 20wt% TaO_x_, which had no significant differences in matrix attenuation between polymer types at the start of the study, Figure 2(a). In non-degrading PCL phantoms, changes in X-ray attenuation at early time points were linked to the constraint of the phantom by surrounding tissues, with X-ray attenuation then leveling off throughout the remainder of the study. In contrast, PLGA matrices had a tendency to experience more severe compaction of the matrix after implantation, leading to complete structural collapse by week 6 for PLGA 50:50, the fastest degrading polymer, Figure 2(a). At week 20, the structural collapse, and densification of the nanoparticles that remained within the implant site led to significantly higher apparent attenuation in areas where the PLGA 50:50 phantoms had been. This was true for phantoms incorporating both 5 and 20 wt% TaO_x_ nanoparticles, Figure 2(b). Densification of nanoparticle contrast agent in PLGA phantoms was further corroborated by increased X-ray attenuation after normalization by the measured volume of matrix present at the implantation site, Figure 2(c). When considering the percentage change in matrix attenuation throughout the 20 week study, Figure 2(d), comparable values were observed in PLGA 50:50 devices with 5wt% TaO_x_ and 20wt% TaO_x_: 204.6 ± 41% and 191 ± 34%, respectively in the intramuscular site.

**Figure 2:**
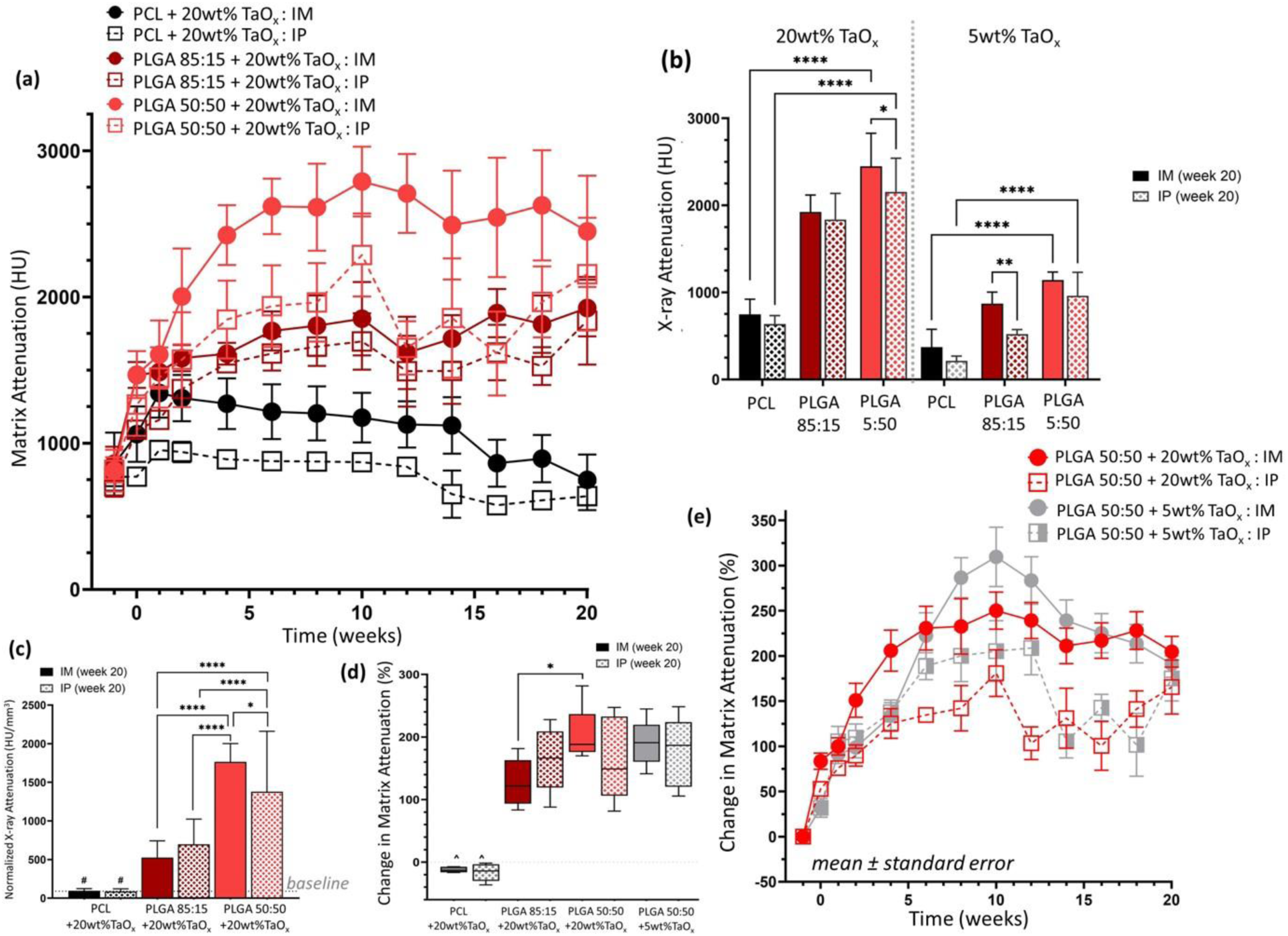
Radiopaque phantoms implanted in vivo could be imaged over 20 weeks. (a) X-ray attenuation of the matrix, in Hounsfield Units (HU), was serially monitored for phantoms with 20wt% TaO_x_ nanoparticles. (b) While initial matrix attenuation was consistent across groups, final X-ray attenuation was dependent on the polymer matrix and implantation site. (c) For non-degrading polymer matrices, X- ray attenuation remained constant when normalized to measured matrix volume, but not for phantoms made from degrading polymers; baseline at day 0 is marked with a dashed line. (d) Degrading phantoms experienced a high percentage change in X-ray attenuation over 20 weeks, which were consistent with both 5 and 20wt% TaO_x_ incorporation, as shown in (e) the change in matrix attenuation of PLGA 50:50 over time, reported as mean ± st error. * p < 0.05, ** p < 0.01, **** p < 0.0001, # significantly different from all PLGA groups (p < 0.01), ^ significantly different from all PLGA groups (p < 0.0001). Unless noted, all graphs show mean ± st deviation.

The kinetics of collapse for fast degrading PLGA 50:50 phantoms were comparable between 20 and 5wt% TaO_x_ nanoparticle addition as well. A peak X-ray attenuation was reached at week 10, Figure 2(e), demonstrating that collapse was mediated by the polymer matrix. During degradation, the molecular weight of the polymer decreases via chain scission prior to complete collapse of mechanical properties [3]. It has been reported that PLGA 50:50 molecular weight degrades with a half-life of less than 2 weeks in vivo, a timeline that fits with the current results [4]. Throughout, PLGA 85:15 change in matrix attenuation fell between non-degrading PCL and PLGA 50:50. This is consistent with a matrix that is less susceptible to hydrolytic degradation than PLGA 50:50 and therefore it does not begin to significantly collapse until after 12 weeks. Previous in vitro studies have shown that PLGA 85:15 does not undergo significant changes in volume or mechanics until at least 16 weeks [21,23,28] and in vivo studies note a half-life of 9-12 weeks depending on device porosity [4,23].

Degradation profiles for the phantoms (Figure 3) were created based on measurements of device volume and structure. Mechanical properties are also an important factor for successful tissue engineering. However, loss of mechanics do not scale to volume or mass loss in polymer systems, but occur significantly faster [21,28]. While PLGA 85:15 might have a half-life of 9-12 weeks in volume, a substantial loss of mechanics has been shown to occur before 16 weeks in vivo [29]. While in vivo monitoring cannot necessarily predict all aspects of device functionality, it can inform the timeline for end-point testing, and hint at mechanical stability by examining when the porous structure is no longer present.

**Figure 3:**
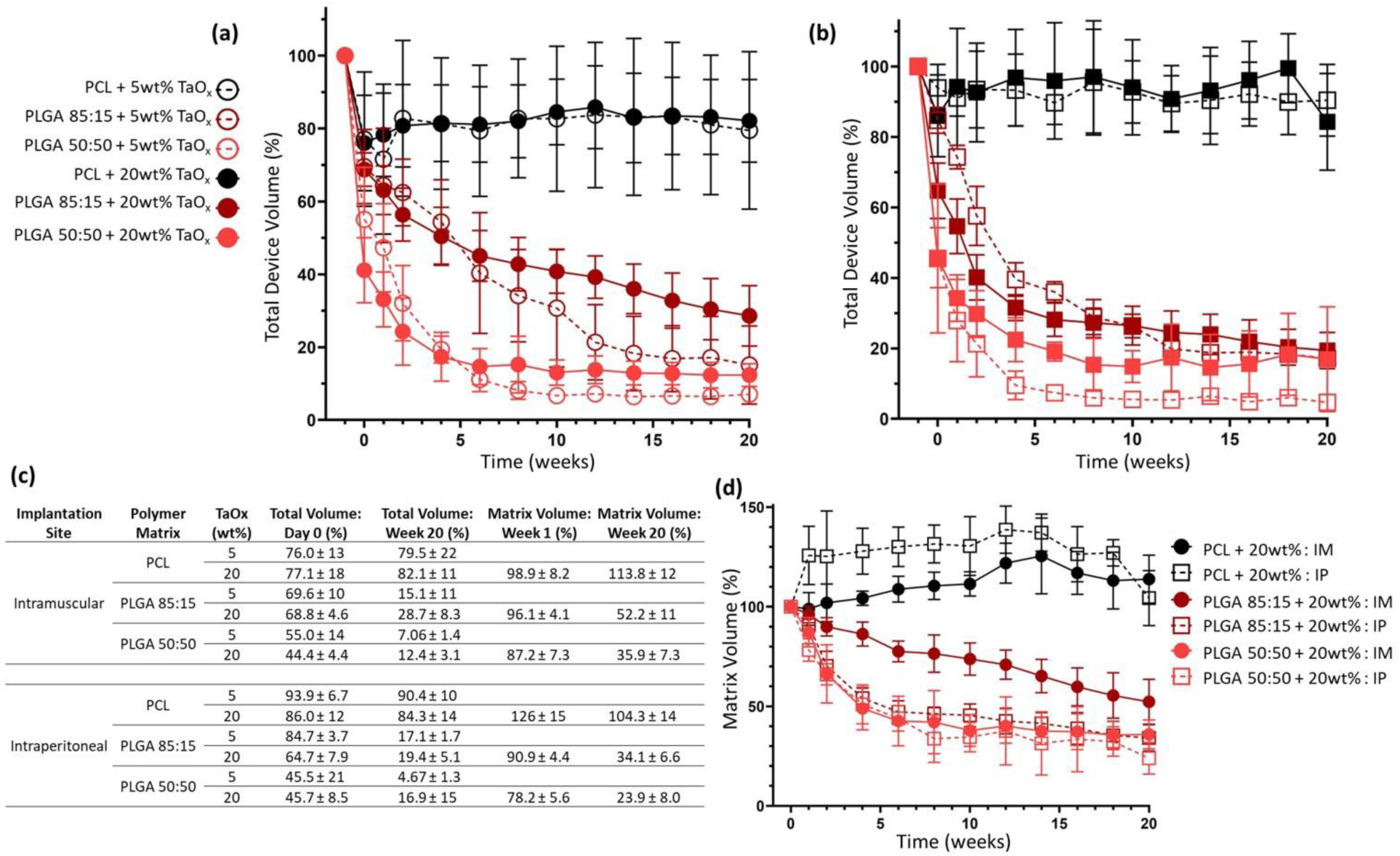
Phantom degradation evaluated in vivo via μCT imaging (groups of n=6). Total volume of phantoms with 5 and 20wt% TaO_x_ was tracked over 20 weeks after (a) intramuscular implantation and (b) intraperitoneal implantation. (c) Significant changes in total volumes and matrix volumes at initial and final time points were observed, and tabulated. (d) Incorporation of 20wt% TaO_x_ nanoparticles allowed quantification of matrix volume in the phantoms; when normalized to post-implant volume swelling and degradation kinetics were highlighted. All data shown as mean ± st deviation.

Understanding and predicting the degradation of polymer devices for clinical use remains challenging even after conducting preclinical studies, as different animal models have differing immune responses and device degradation profiles [30]. Also, using fluorescently tagged hydrogels, it has been demonstrated that differences in degradation rates can be linked to morphological traits of the implanted device, like surface area [31]. Previous literature has utilized X-ray tomography to monitor degradation in vivo, successfully tracking iodinated polyethylene glycol (PEG) based hydrogels that had for up to 30 days in various injection sites [14,32]. During these studies, like the present case, phantoms have increased contrast over time due to compaction of the phantom or contrast densification [14].

While hydrogels are useful for delivering drugs or cells, many implanted devices must mechanically stabilize an injury site and remain in place until new tissue can be formed. This requires longer term monitoring and a different set of polymeric materials. Chemical modification, such as iodination, has been applied to polyesters like PLGA and PCL, but less attention has been paid to their degradation profiles and it remains uncertain if the iodinated groups remain associated with the polymer structure for the long term [12,33]. Utilizing nanoparticles as a contrast agent allows the independent tuning of both the contrast agent and polymer matrix. Further, nanoparticles based on different chemistries, such as gold and iodine, facilitate the differentiation of various components in a tissue engineered system via emerging color-CT technologies, allowing cell and materials components to be individually tracked in vivo [9]. In this study, we have successfully tracked in vivo phantom response over 20weeks, highlighting when structural collapse occurs, signaling loss of mechanical stability. This was affected by both polymer composition and implantation site.

### 2.2 Effect of Composition

In the field of tissue engineering, polymer degradation is supposed to match tissue regeneration, ensuring that device is replaced by functioning tissue over time. This is believed to be a key reason for implanted biomedical device failures, and drives the search for materials with novel and controlled properties. As the degradation rate required is not tied to a particular timeline, but can vary with the biological application, a variety of polymer types were used to simulate tissue engineering devices in the present study, from virtually non-degrading (PCL) to very fast degrading (PLGA 50:50). As the the polymer matrix made up the bulk of the phantom composite materials, degradation profiles were dominated by the polymer hydrolysis rate [21].

Over 20 weeks, phantom volume was measured as a surrogate for device degradation, Figure 3(a-b). As expected, PCL phantoms had no significant change in volume over 20 weeks, after the initial implantation, while PLGA 50:50 phantoms experienced rapid volume decreases. PLGA 85:15, a mid- degrading polymer, was in between the two extremes, a trend visible regardless of the amount of TaO_x_ nanoparticles incorporated, in both the intramuscular and intraperitoneal spaces. After 20 weeks in the intramuscular space, PCL phantoms had retained >79% of their original volume, while PLGA phantoms had < 30% remaining. However, with higher TaO_x_ addition, the final apparent volume was affected, with 20wt% TaO_x_ showing significantly higher post-degradation volumes than phantoms with 5wt% TaO_x_ nanoparticles, Figure 3(c).

Volumes have been utilized as surrogates for device degradation in the past, with the assumption that there is a matched excretion rate of the contrast agent compared to the matrix polymer [10]. When the material components making up the phantom composition are on the extremes of diffusion, this assumption can break down at the end points of degradation. This was observed with PLGA phantoms, where the excretion rate of TaO_x_ nanoparticles from tissues is slower than the diffusion of oligomer and monomer left over from polymer chain scission, as observed in subcutaneous degradation studies [27]. While the hydrophobic polymer matrix remains intact, the nanoparticles remain associated with it [21].

However, after device structure has been lost, following significant scission of the polymer chains [3], it isn’t known if the hydrophobic nanoparticles are internalized via macrophages present at the site, or become associated with hydrophobic elements in the ECM, slowing their removal. Additional steps may be required to match the excretion of nanoparticles to fast-degrading polymer matrices by altering the size of the nanoparticles or increasing the hydrophilicity [34]. While chemically binding contrast agent to the polymer chain may be a more direct method of coupling degradation of contrast agent and polymer, the addition of iodinated contrast agents can affect measured the degradation rates of phantoms, therefore altering biological response [14]. During in vitro experiments, the decoupled rates of diffusion are not apparent, given the lack of tissue matrix around phantoms. In vivo, while the end point degradation kinetics of oligomers cannot be tracked utilizing the current TaO_x_ nanoparticles, this study still provides a compelling insight into the kinetics of phantom collapse and allows comparison across multiple implant sites.

### 2.3 Effect of Implantation Site

In determining in vivo degradation profiles for biomedical devices, a variety of implantation sites have been used, including subcutaneous, intramuscular and intraperitoneal. Each site contains a unique combination of immune cells, tissue types, hormones and mechanics. This complex biological milieu has a significant effect on the degradation of biomaterials [31,35]. For devices designed for specific applications, testing is generally carried out in a site related to the final purpose. However, where general materials properties are the focus of investigation, there is no standardized protocol in the literature, leading to difficulty comparing various studies of in vivo polymer degradation. Therefore, the current study separated out effects due to composition changes and those due to biological factors related to the implant site.

Two different implantation sites were investigated: the intraperitoneal site (Figure 4) and the intramuscular site (Figure 5). It was found that implantation site had a significant effect on degradation, but the severity of the effect was linked to the polymer matrix properties. In all cases, surgical implantation created severe initial reductions in phantom volume compared to pre-implantation volumes, Figure 3(c), as has been noted in literature [27]. This was most severe for PLGA 50:50 with 20wt% TaO_x_ nanoparticles, where total volume dropped to 44.4 ± 4.4% and 45.7± 8.5% of the pre-implantation volume when placed intramuscularly or intraperitoneally, respectively. Initial volume reduction was not as severe in PLGA 85:15 matrices where post-implantation volume was 68.8±4.6 and 64.7±7.9% and even less severe for PCL, which retained close to 80% of pre-implantation volume.

**Figure 4:**
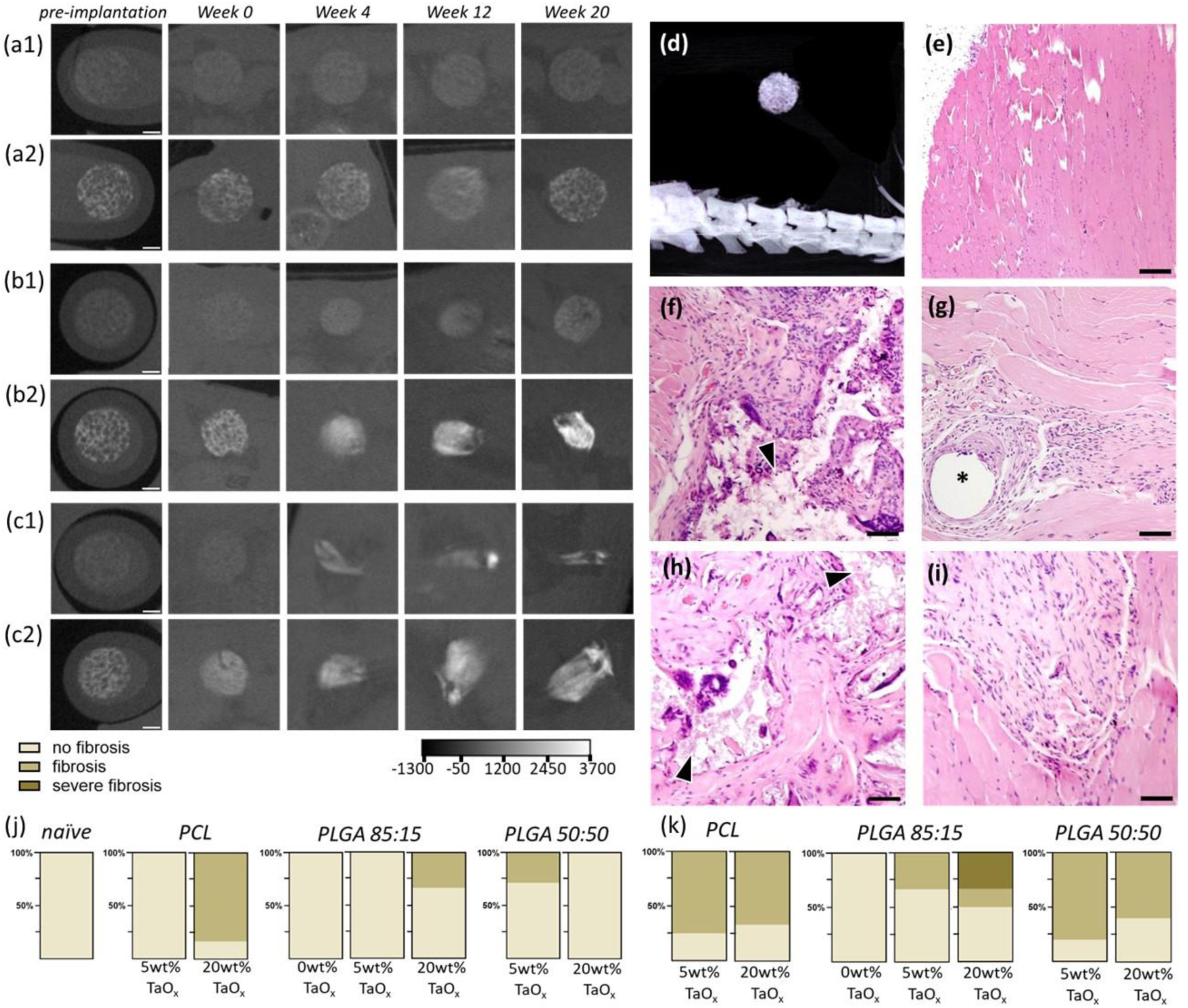
Radiopaque phantoms implanted intraperitoneally (n=6 per group) degraded over time. (a-c) In situ monitoring via μCT was performed over 20 weeks to follow degradation kinetics for (a) PCL phantoms, (b) PLGA 85:15 phantoms, and (c) PLGA 50:50 phantoms incorporating (1) 5wt% TaO_x_ nanoparticles or (2) 20wt% TaO_x_ nanoparticles; the HU scale is consistent across all images. (d) A 3D rendering of the intraperitoneal implantation site shows the placement of phantoms above the spine.

**Figure 5:**
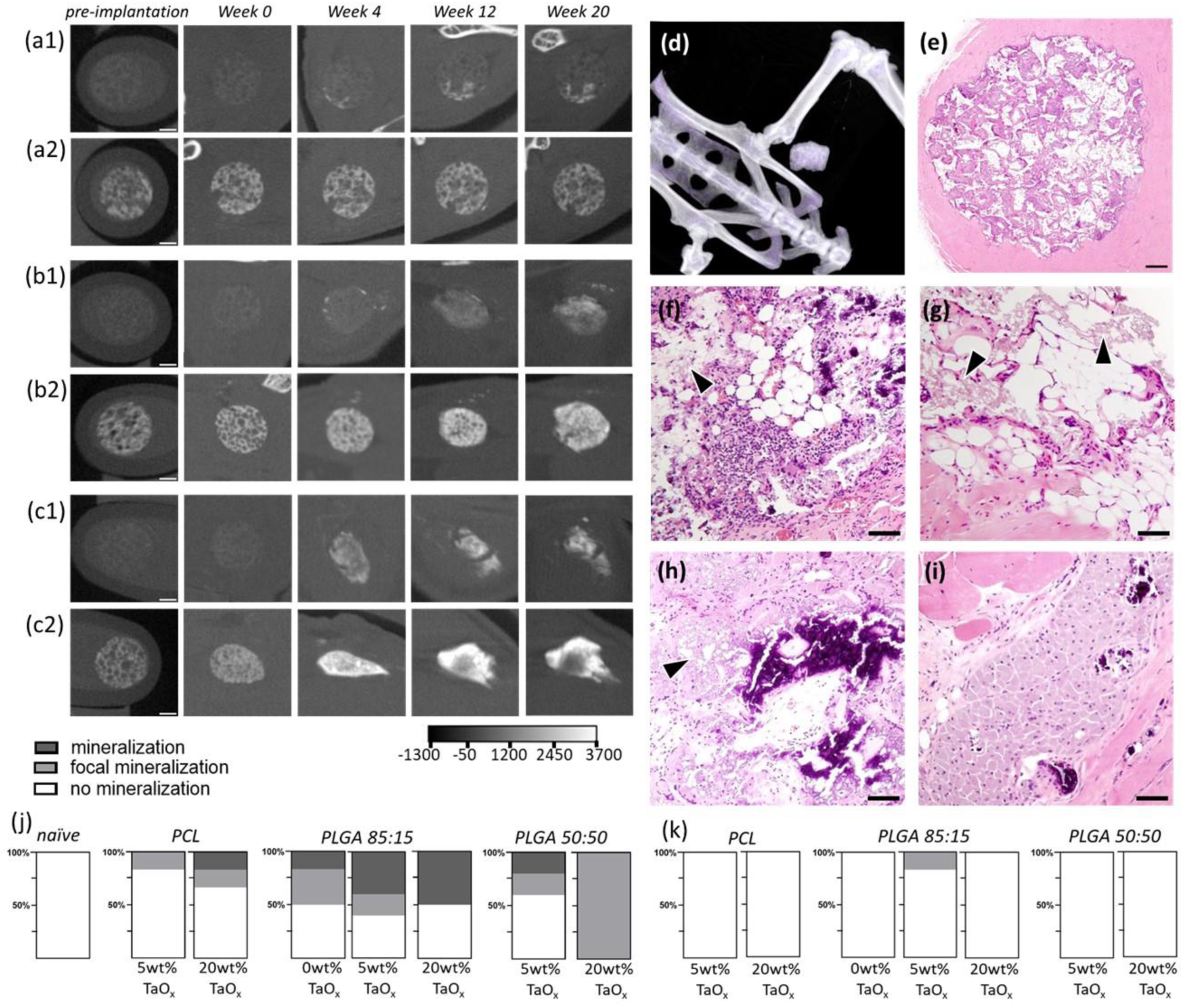
Radiopaque phantoms implanted intramuscularly (n=6 per group) were prone to mineralization. (a-c) In situ monitoring via μCT followed degradation over 20 weeks of (a) PCL phantoms, (b) PLGA 85:15 phantoms, and (c) PLGA 50:50 phantoms incorporating (1) 5wt% TaO_x_ nanoparticles or (2) 20wt% TaO_x_ nanoparticles The presence of mineralized areas was visible was bright specular regions in the phantoms with 5wt% TaO_x_. The HU scale is consistent through all images. (d) A 3D rendering of the intramuscular implantation site shows the placement of phantoms, that were (e) well integrated into muscle tissue after 20 weeks, shown via H&E staining of a PCL phantom containing 5wt% TaO_x_. At higher magnification, the integration of tissue and formation of mineralized areas was visible in (f) PLGA 85:15 with 0wt% TaO_x_, (g) PCL with 20wt% TaO_x_, (h) PLGA 85:15 with 20wt% TaO_x_ and (i) PLGA 50:50 with 20wt% TaO_x_. Mineralization occurred significantly in the (j) intramuscular site compared to the (k) the intraperitoneal site. The incidence of mineralization increased with greater TaO_x_ nanoparticle content. Scale bar: (a-c) 1 mm, (e) 300 μm (f-i) 15μm. Arrowheads: indicate remaining polymer matrix

Measurements of the matrix volume, carried out for phantoms with 20wt% TaO_x_, revealed that while total volume of PCL phantoms might not be significantly altered by implantation site, significant swelling of the matrix occurred at the intraperitoneal site at initial time points. This correlated to a matrix volume of 126 ± 15% of initial post-implantation matrix volume in the intraperitoneal space compared to 98.9 ± 8.2% intramuscularly. The greater mechanical confinement of phantoms at the intramuscular site was further shown with a tendency for increased x-ray attenuation, corresponding to greater physical nanoparticle compaction, Figure 2.

The most significant effects on degradation rate with implantation site occurred for mid-degrading polymer phantoms, those with matrices of PLGA 85:15. This same effect of implantation site on degradation kinetics was not observed in slow- (PCL) or fast- (PLGA 50:50) degrading polymers. In the case of slow-degrading PCL matrices, an effect may be observed at later stages of degradation (> 2 years). For fast degrading PLGA 50:50, the rate of polymer chain scission was not affected by the implant site, leading to compaction of all devices within 8 weeks.

At the intramuscular site, PLGA 85:15 phantoms degraded linearly over time up to week 14 (R^2^ > 0.91). The degradation curve is likely controlled by a combination of polymer chain scission and subsequent diffusion of oligomers and nanoparticles from the site. Since chain length decreases before a measurable effect on mass loss is observed [28], it is likely that removal of oligomers and nanoparticles at the implantation site was a rate limiting step. Indeed, total volume of the phantoms changed more rapidly than matrix volume, indicating that phantom compaction occurs prior to matrix removal, as observed in vitro [21]. Also, the measured degradation kinetics were slower for phantoms incorporating 20wt% TaO_x_ nanoparticles, supporting the idea that measured volume changes are limited by slow nanoparticle diffusion. In contrast to the intramuscular site, PLGA 85:15 phantoms implanted intraperitoneally degraded much faster. The degradation kinetics could be approximated by a second order polynomial fit (R^2^ > 0.95), and mimicked the degradation profiles of PLGA 50:50 phantoms.

Measurable differences in diffusion pathways and drug delivery kinetics have been noted between the intramuscular site and intraperitoneal sites. The intramuscular site represents a highly vascularized tissue bed that has access to systemic circulation without passing first through any organ that could metabolize molecules, such as the liver. In contrast, the peritoneal site offers access to both the circulatory system, with its high surface area and large blood supply, but also to the lymphatic system [36]. Indeed, large molecules are taken up via lymphatics in the intraperitoneal space prior to passing through venous circulation [36]. Studies on bioavailability of large molecule drugs report that concentrations of drug in the bloodstream are highest after intravenous injection, followed by the intraperitoneal route, then the intramuscular route and finally by subcutaneous injection [36].

In terms of phantom degradation, the direct link to the lymphatic system in the intraperitoneal space and a pool of immune cells, may increase degradation kinetics, particularly as PLGA is known to be degraded faster when in contact with inflammatory macrophages [27]. Difference in the kinetics of degradation may also be linked to how the circulatory system is accessed at each site. While there is a large blood supply within the intramuscular space, there are likely more barriers to diffusion related to the ECM proteins around the muscle fibers [37]. In addition, polymers like PLGA that degrade via hydrolyzation are affected by the ability of water to diffuse into the polymer matrix. Where material thickness is constrained, in thin films or in the struts of porous devices, degradation has been found to follow first order kinetics, with glycolic acid units degrading faster [38]. Buildup of acidic byproducts of polymer degradation might also drive increased phantom degradation, but it is not likely the cause of the implant site differences observed here, as peritoneal fluid is buffered around pH 7.5-8 [36]. However, the intraperitoneal space does contain a greater fluid volume than the intramuscular site, and fluid volume has been correlated to increased degradation rates for hydrogel systems [6,31]. Studies have also shown that intraperitoneal spaces have greater immune responses around implanted devices than after subcutaneous implantation, but that placement near adipose tissue triggers the greatest inflammatory responses [35].

Regardless of placement, all phantoms were well tolerated by the body, consistent with the use of these polymers as biomedical devices [39,40]. As expected, surgical implantation of phantoms produced a measurable immune response in vivo, demonstrated by granulomatous cellulitis around and within phantoms. This was consistent regardless of implantation site, with a tendency for PCL phantoms to have a more severe granulomatous cellulitis, likely due to the fact that the polymer remains intact over the 20-week period, Supplemental Figure S3. In other phantom matrices, the inflammation had moved to a more chronic stage associated with tissue repair. The severity of the inflammatory response was not due solely to nanoparticle addition and did not scale with the amount of TaO_x_ nanoparticles incorporated.

Throughout the study, CT scanning was conducted every two weeks to minimize radiation exposure, as this timing has been shown not to dampen the systemic immune system of mice [27]. However, as the physiological result of ionizing radiation due to localized scans is dependent on the sensitivity of surrounding tissues [41], white blood cell count was taken at 20 weeks to monitor any systemic changes in the immune response. There were no significant changes in white blood cell count related to X-ray exposure in the study (Supplemental Table S2). Further, biological responses around phantoms that were not monitored via μCT were not significantly affected compared to phantoms that were imaged.

After 20 weeks, tissues were collected and stained to determine the inflammation and fibrosis present and compared to (e) naive tissue: (f) PCL with 5wt% TaO_x_, (g) PLGA 50:50 with 5wt% TaO_x_, (h) PCL with 20wt% TaO_x_ and (i) PLGA 50:50 with 20wt% TaO_x_. The incidence of fibrosis was generally lower in the (j) intramuscular site than (k) the intraperitoneal site, but fibrosis was not mediated by TaO_x_ nanoparticle content or polymer type. Scale bar: (a-c) 1 mm, (e-i) 15μm. Arrowheads: indicate remaining polymer matrix; * suture material.

The intraperitoneal space is known to have a high component of immune cells, and the ability to allow diffusion of proteins and molecules into systemic circulation or directly to the lymphatic system [36]. In the intraperitoneal space, phantoms of PLGA were compacted over time, with gradual loss of a distinct porous morphology, Figure 4(a-c). Histology showed the presence of many immune cells surrounding intraperitoneal phantoms, Figure 4(f-g). The immune response had already subsided for phantoms of fast-degrading polymer (PLGA 50:50) compared to non-degrading PCL. The immune cells remaining after PLGA 50:50 degradation could be associated with residual nanoparticles, or with the non-degrading suture used to secure the implants and mark the site of implantation post-degradation. In μCT scans, the presence of this suture at the site was noted as a dark spot within the bright signal volume. In general, fibrosis was observed with higher frequency for devices implanted in the intraperitoneal space compared to an intramuscular site, Figure 4(j-k). However, the fibrosis present was generally not severe and there were no consistent trends in incidence of fibrosis with increasing TaO_x_ content or between different polymer matrices.

Compared to the intraperitoneal space, physical degradation of phantoms in the intramuscular space, and by extension biological response, was expected to be modulated by mechanical forces acting at the site. In addition to mechanical forces, skeletal muscle is well innervated and has innate immune cells and the ability to recruit additional cells post-injury [42]. The intramuscular space constrained degrading phantoms, causing collapse and elongation over time, Figure 5(a-c). After 20 weeks of implantation, phantoms were well infiltrated with tissue throughout their structure, Figure 5(e). In addition, mineralization was observed within the porous phantom structure. These areas were visible in μCT scans as regions of increasing X-ray attenuation over time, particularly in phantoms incorporating 5wt% TaO_x_ nanoparticles, Fig 5(a-c). Some incidence of mineralization was present in all types of polymer matrices, Fig 5(f-i), with the presence of TaO_x_ nanoparticles increasing the incidence and severity of mineralization responses. Mineralization did not occur significantly in the intraperitoneal space, Figure 5(j-k).

It is known that tissue engineering devices incorporating ceramic nanoparticles, particularly those containing calcium phosphates can induce bone formation when implanted intramuscularly [43]. In literature, this is a property tied to porous devices, that also exhibit a rough surface morphology, and has also been linked to a higher potential for bone formation when the same devices are placed in bone defects [43]. The radiopaque phantoms used in the current study fit into this description, having nanoscale roughness that increases with the incorporation of nanoparticles into polymer matrices [25]. While most studies on osteoinduction concentrate on calcium phosphate addition, as the key mineral in bone, bone repair has been shown in the past on composites of polyetheretherketone (PEEK) and tantalum nanoparticles, increasing with tantalum addition [44].

The foreign body reaction to tissue engineering constructs is a multi-step process, beginning with an acute inflammatory phase that generally subsides within a week into a more sustained foreign body response local to the implant interface [45]. The action of inflammatory cells is followed by fibroblast invasion and revascularization. Depending on the environmental cues, this could result in fibrous encapsulation of the implant that can impair healing. Not only environmental cues, but interactions with the implanted material can also direct the biological response. In vivo, it has been observed that PCL and PLGA matrices trigger different immune responses, with PCL leading to higher rates of revascularization [46]. Some of the differences in biological responses to polymer matrices are linked to the degradation products of those polymers, such as lactic acid, which can affect cellular metabolic activity [26,47].

Introduction of contrast agents into polymers can also trigger increased or chronic inflammatory responses, such as the incorporation of iodinated groups into PCL that triggered chronic inflammatory responses and greater fibrous encapsulation [48]. Previous in vitro studies have shown that alterations in biological response with incorporation of TaO_x_ nanoparticles into phantoms is mediated by changes to phantom properties, such as increased surface nanotopography [25]. Surface topography can affect foreign body reaction via protein interactions and cellular adhesion, but many factors modulate biological response making it difficult to predict a priori [45,49] The site specific biological interactions observed in the current study, included mineralization in muscle tissue and greater incidence of fibrosis intraperitoneally.

The excretion of contrast agents after device degradation remains a concern for determining the overall toxicity of implanted phantoms. The toxicity, clearance and excretion pathway of nanoparticles injected into blood circulation has been closely linked to their size and chemistry [50]. For tantalum nanoparticles < 4 nm diameter administered intravenously, which have been proposed as clinical contrast agents, there is rapid clearance via the kidneys, liver and spleen [51]. With increasing size, nanoparticles tend to have shorter retention in the blood circulation and are excreted less by the kidney and more in liver and spleen [16,50]. Nanoparticle chemistry can also play a role in clearance and excretion pathways, offering ways to tune the clearance and overall toxicity to the nanoparticle contrast agent [51].

When nanoparticles are incorporated into tissue engineering devices, both nanoparticles and polymer degradation products must first diffuse through tissue and the circulatory system prior to being removed via secretory organs. While size often dominates discussions on nanoparticle toxicity, affecting cellular interactions, it also determines nanoparticle movement through the dense ECM of collagen bundles [52]. It has been shown that hydrophobic nanoparticles, such as those used in the current study, are further hindered in diffusion due to hydrophobic interactions with the ECM matrix [34]. Together this accounts for the decoupling of the polymer and nanoparticle diffusion observed at late time points of phantom degradation in the present study.

In situ monitoring highlights the interactions between devices and the biological environment and how these are specific for both device composition and placement. Despite challenges of final nanoparticle clearance that have yet to be overcome, there remains a great deal to explore using radiopaque phantoms. Not only can materials properties like degradation be monitored, but tissue and implant movement can be tracked non-invasively to extend our understanding of mechanical forces during physiological movement and growth, Figure 6. Thus, in situ monitoring is poised to facilitate clinical translation of biomedical devices and ensure patients reap the benefits of tissue engineering research.

**Figure 6:**
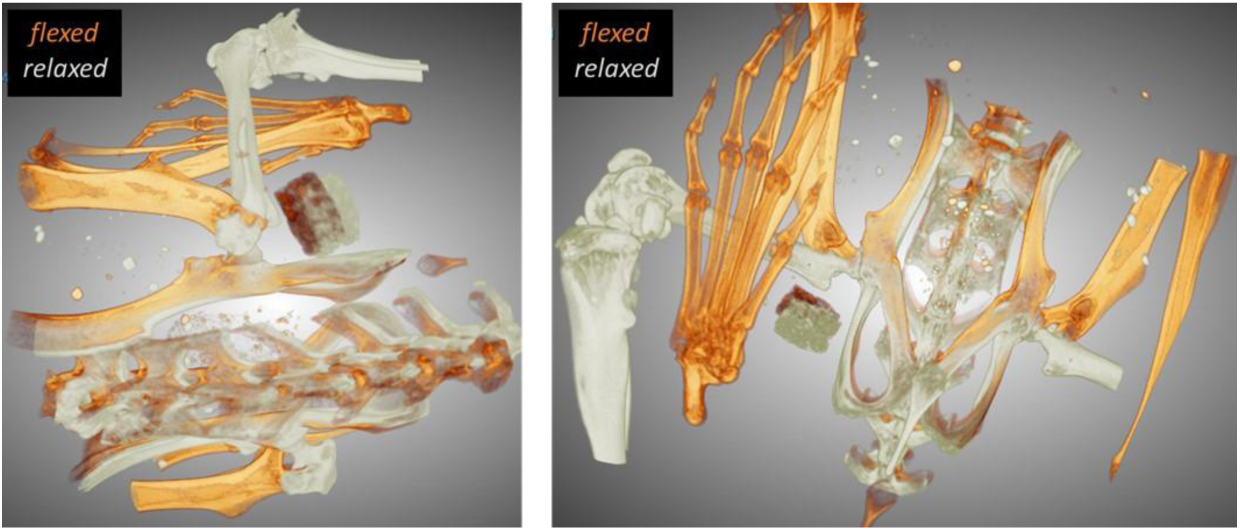
Radiopaque phantoms can facilitate in situ monitoring of materials properties, like degradation, and also physiological forces during movement, as shown in 3D renderings of radiopaque phantoms positioned intramuscularly, overlayed between a relaxed and flexed leg position in different views.

## 3 Conclusion

In situ monitoring of implanted medical devices has been identified as a growing concern, both to better inform tissue engineering device design and to monitor devices in the clinic to improve patient outcomes. For polymeric implanted devices, biological monitoring is problematic due to poor contrast in commonly used clinical imaging techniques, necessitating the use of contrast agents. With the introduction of a radiopaque TaO_x_ nanoparticle into polymer matrices, phantoms that mimicked tissue engineering devices were created and in vivo degradation profiles were successfully monitored via CT. Long-term monitoring over 20 weeks demonstrated sensitivity to phantom volumetric changes due to swelling and compaction post-implantation. Phantom degradation was driven by the polymer matrix properties, with the amount of nanoparticle contrast agent incorporated having little influence on overall kinetics. Importantly, given the discrepancies in published literature on polymer degradation, implantation site of the phantom was found to have a significant effect on degradation kinetics, particularly for PLGA 85:15 that has a degradation profile over months. Due to differences in inflammatory cells and fluid flow present, not only degradation but the foreign body response was also determined by the implantation site, with intramuscular implantation stimulating the formation of mineralized tissue while intraperitoneal phantoms had higher incidences of fibrosis. After phantom collapse, it was observed that the diffusion of nanoparticles from the implantation site occurred more slowly than the excretion of the corresponding polymer, highlighting the need for targeted chemistry of the contrast agent component to ensure better clearance from tissues. Overall, this study shows the effectiveness of long-term monitoring for understanding materials properties and enabling the translation of tissue engineering from the bench to the bedside.

## 4 Materials & Methods

The study utilized three types of biocompatible polymers: polycaprolactone (PCL), poly(lactide-co-glycolide) (PLGA) 50:50 and PLGA 85:15. PCL (Sigma Aldrich) had a molecular weight average of 80 kDa. PLGA 50:50 (Lactel/Evonik B6010-4) and PLGA 85:15 (Expansorb® DLG 85-7E, Merck) were both ester terminated and had a weight average molecular weight of between 80-90 kDa.

Hydrophobic nanoparticles of amorphous TaO_x_ were manufactured based on our previous procedure with one minor change [19]. Hydrophobicity is imparted by coating nanoparticles with hexadecyltriethoxysilane (HDTES, Gelest Inc., cat no SIH5922.0), an aliphatic organosilane. The coating has been shown to produce polymer composites with homogeneous nanoparticle distributions [21]. The spherical particles had diameters in the range of 3-9 nm (Supporting Information S1).

### 4.1 Composite manufacture of phantoms

All polymers used had molecular weights between 80-90 kDa. Polymers were solubilized in suspensions containing TaO_x_ nanoparticles in dichloromethane (DCM, Sigma); PCL was used at 8 wt %, while PLGA was used at 12 wt%. The amount of TaO_x_ (0-20wt%) was calculated based on the weight percent of the total dry mass (polymer + nanoparticle). To introduce microporosity to the phantom walls, sucrose (mean particle size 31 ± 30 μm, Meijer) was added to the solution, so that the final slurry was 70 vol % sucrose and 30 vol % polymer+nanoparticles. This was followed by addition of NaCl (Jade Scientific) at 60 vol% of the total polymer + nanoparticle volume to create macroporosity. The suspension was vortexed for 10 minutes and pressed into a silicon mold that was 4.7 mm diameter, 2 mm high. After air drying, phantoms were removed and cut down to 3mm diameter using a biopsy punch. Finally, phantoms were washed for 2 hours in distilled water, changing the water every 30 minutes to remove sucrose and NaCl. After air drying phantoms overnight, they were stored in a desiccator prior to use.

Nominal nanoparticle concentration was measured by thermogravimetric analysis (TGA) and the associated radiopacity was quantified via micro-computed tomography (μCT). The information is reported in supplemental information (Sections S.2-3). In addition, TGA was used to confirm the complete removal of sucrose and salt with the washing steps described above. No inhomogeneity in phantom microstructure or radiopacity was encountered to suggest sedimentation of nanoparticles or incomplete mixing of suspensions.

### 4.2 Microscopy

Scanning electron microscopy (SEM) images were taken of phantoms in the dry state. To prepare for SEM, phantom cross-sections were adhered to 13 mm aluminum stubs and sputter coated with gold (approx 30 nm thickness) in an Emscope Sputter Coater model SC 500 (Ashford, Kent, England) purged with argon gas. Microstructure was examined in a JEOL 6610LV (tungsten hairpin emitter) scanning electron microscope (JEOL Ltd., Tokyo, Japan) at 12 kV and a working distance of 12 mm.

### 4.3 Micro-computed tomography (µCT)

All tomography images were obtained using a Perkin-Elmer Quantum GX. Prior to implantation, devices were imaged at 90 keV, 88 µA. In vitro imaging was conducted with a 25 mm field of view at a 50 µm resolution. After acquisition, individual devices were sub-reconstructed using the Quantum GX software to 12 µm resolution. Hydrated phantoms used for serial monitoring were imaged 24 hrs before implantation.

In vivo μCT on mice was performed at 25 mm field of view (4 min total scan time) at 50 μm resolution, centered on the implanted phantom. During acquisition, mice were anesthetized using an inhalant anesthetic of 1–3% Isoflurane in 1 L min^−1^ oxygen. Mice (n = 72) were scanned 24 hours post-implantation, on week 1, and then every other week until week 20 was reached (week 1, 2, 4, 6, 8, 10, 12, 14, 16, 18, 20). Mice (n = 3) were kept as naive controls and an additional cohort of mice (n = 12) received surgical implants but were not scanned in μCT until after euthanasia at week 20. Total cumulative radiation dosage was 35.4 Gy over 20 weeks and was delivered to local regions around the phantoms. The physiological result of radiation due to localized scans is dependent on the sensitivity of surrounding tissues such as adipose tissue or bone marrow [41]. In the present study, this was tracked via complete blood count following euthanasia (Table S2, Supplemental). Scans that were performed at week 20 took place immediately following euthanasia (n = 84). After all acquisitions, individual phantoms were again sub-reconstructed using the Quantum GX software to 12 µm resolution.

#### 4.3.1 Tomography analysis

To analyze 12 µm sub-reconstructions tomography scans of phantoms incorporating 20wt% TaO_x_ nanoparticles, we developed a custom process to analyze the volumetric µCT images in order to extract in a reproducible manner summary measures of the scaffold properties. This tool, developed in MATLAB (v R2021b, Mathworks, Natick, MA), first performed segmentation using a Ostu-based threshold technique to identify the most attenuating voxels in the image [53]. These segmented voxels went through a series of volumetric morphologic operations (erosion and dilation) to identify connected voxels of at least 1 mm^3^ in size. Finally, we calculated measures from these regions to summarize the number of pores inside the segmented envelope, pore volume, total attenuation, and total volume.

For phantoms with 5wt% TaO_x_, scans at 50 μm resolution were used as reliable segmentation from background tissue was difficult. Prior to collapse of the structure, gross features of the phantoms (thickness, diameter) were analyzed using Image J; this corresponded to 0-20 weeks for PCL phantoms, 0-10 weeks for PLGA 85:15 phantoms and 0-2 weeks for PLGA 50:50 phantoms. In Image J, image stacks were opened and rotated, using built in functions, to measure thickness and diameter in 5 planes, which were averaged. From the diameter and thickness, a “gross volume” was defined as the volume occupied by a solid cylinder with the corresponding thickness and diameter. After collapse of the internal macroporosity in the phantoms, analysis of phantom volume was performed using ITK SNAP [54]. The polymer phantom was segmented from the background tissue and the volume of the matrix and its average intensity was calculated. Regardless of the program used for calculation, the gross volume of phantoms consisted of both the polymer matrix volume and the volume of the pores. Three-dimensional reconstructions from in vivo imaging were made using Dragonfly (v.2022.2) built in functionality.

### 4.4 In vivo study

#### Phantom preparation

Phantoms incorporating 0-20 wt% TaO_x_ nanoparticles were produced as detailed above. After preparation of phantoms (3 mm diameter, 2 mm thickness) were soaked in 70% ethanol for 30 min. The ethanol was replaced by sterile PBS and phantoms were centrifuged for 10 minutes at 11,000 rpm. The PBS was replaced and the phantoms were left in sterile PBS until implantation.

#### Surgical implantation

All procedures were performed in accordance with IACUC-approved protocols and Veterinary guidelines at Michigan State University. BALB/c Mice (n = 87 adult males, 2 months old; Charles River Laboratories) were used for this surgical implantation and μCT imaging study. Mice (n = 84) were surgically implanted with an assigned phantom consisting of varying material and differing weight percent of TaO_x_ nanoparticles. The phantom was installed within a single implantation surgical site located at either the intramuscular layer of the right leg or the intraperitoneal space on the ventral abdomen, Figure 1. Each group consisted of 6 individuals. Mice (n = 3) were kept as naive controls. Each animal implanted was administered analgesia at least 20 min prior to making the initial incision, including prophylactic Ampicillin (25 mg kg−1; SID) administered S.C., Meloxicam (5 mg kg−1; SID) administered S.C. in the right dorsal lateral flank of the animal, and local infiltration of 2% Lidocaine (diluted to 0.5%) was administered S.C. along the intended incision site just prior to making the cutaneous incision (7 mg kg−1 Max dose; SID). Animals were anesthetized via inhalant isoflurane (3–4% isoflurane in 0.8–1 LPM oxygen for induction) and maintained via inhalant isoflurane during surgery (1– 3% isoflurane in 0.8-1 LPM oxygen). Supplemental heat was provided via recirculating warm water blankets during anesthesia induction, patient preparation, surgery, and patient recovery. Each phantom was sutured to surrounding tissue with at least one single interrupted suture using 9-0 PROLENE (Polypropylene) monofilament Suture for in situ location retention. Incision sites were closed in multiple layers where necessary using 5-0 COATED VICRYL (polyglactin 910) Suture and cutaneous layers were closed using 5-0 PDS-II (polydioxanone) Suture. Following animal recovery, Meloxicam (5 mg kg−1; SID) and Buprenorphine (2 mg kg−1, BID, every 8–12 h) were administered S.C. in the left or right dorsal lateral flank of the animal for 48 h following surgery. Post-operative clinical observation, body weight assessment, and health score assessment were performed for 7–14 days postoperatively.

#### End of study

Euthanasia was performed on week 20 (n = 87). At termination, animals were anesthetized via inhalant isoflurane (3–4% isoflurane in 0.8–1 LPM oxygen for induction) and sacrificed via CO_2_ overdose asphyxiation. Following euthanasia, blood samples were collected for complete blood count (Table S2, Supplemental Information). Tissue samples of the heart, brain, kidney, liver, spleen, and the phantom and surrounding tissue were also collected post-euthanasia. Tissues were fixed in 10% neutral buffered formalin, then embedded in paraffin for sectioning and stained with hematoxylin and eosin (H&E). Stained slides were used for histopathological assessment by a certified veterinary pathologist, examining signs of inflammation and fibrosis.

### 4.5 Statistics

Statistics were performed using GraphPad 10.3.1. Data was analyzed via ANOVA, followed by Fishers LSD test. In all cases, α < 0.05 was considered significant, with a 95% confidence interval. Unless noted, all data is reported as mean ± standard deviation.

## Acknowledgments

The authors would like to thank the Center for Advanced Microscopy (Michigan State University) for obtaining electron microscopy images and the Veterinary Diagnostic Laboratory (MSU) for histological staining. We also thank Ethan Tu and Adam Alessio for contributing the analysis code for evaluating CT images. This study was supported by the National Institute of Biomedical Imaging and Bioengineering of the NIH under award number R01EB029418. The content is solely the responsibility of the authors and does not necessarily represent the official views of the National Institutes of Health.

## References

[1] M. Veletic, E. H. Apu, M. Simic, J. Bergsland, I. Balasingham, C. H. Contag, and N. Ashammakhi, “Implants with sensing capabilities,” Chemical Reviews 122, pp. 16329–16363, 2022.

[2] C. Gil, M. L. Tomov, A. S. Theus, A. Cetnar, M. Mahmoudi, and V. Serpooshan, “In vivo tracking of tissue engineered constructs,” Micromachines 10, p. 474, 2019.

[3] F. Alexis, “Factors affecting the degradation and drug-release mechanism of poly(lactic acid) and poly[(lactic acid)-co-(glycolic acid)],” Polymer International 54, pp. 36–46, 2005.

[4] L. Lu, S. J. Peter, M. D. Lyman, H.-L. Lai, S. M. Leite, J. A. Tamada, S. Uyama, J. P. Vacanti, R. Langer, and A. G. Mikos, “In vitro and in vivo degrdation of porous poly(dl-lactic-co-glycolic acid) foams,” Biomaterials 21, pp. 1837–1845, 2000.

[5] K. Pawelec, S. Chakravarthy, J. Hix, K. Perry, L. van Holsbeeck, R. Fajardo, and E. Shapiro, “Design consideration to facilitate clinical radiological evaluation of implantable biomedical structures,” ACS Biomaterials Science & Engineering 7(2), pp. 718–726, 2021.

[6] W. Wang, J. Liu, C. Li, J. Zhang, J. Liu, A. Dong, and D. Kong, “Real-time and non-invasive fluorescence tracking of in vivo degradation of the thermosensitive pegylated polyester hydrogel,” J. Mater. Chem. B 2, p. 4185, 2014.

[7] A. Weems, J. Szafron, A. Easley, S. Herting, J. Smolen, and D. Maitland, “Shape memory polymers with enhanced visibility for magnetic resonance- and x-ray imaging modalities,” Acta Biomaterialia 54, pp. 45– 57, 2017.

[8] T. A. Finamore, T. E. Curtis, J. V. Tedesco, K. Grandfield, and R. K. Roeder, “Nondestructive, longitudinal measurement of collagen scaffold degradation using computed tomography and gold nanoparticles,” Nanoscale 11, p. 4345, 2019.

[9] E. Cuccione, P. Chhour, S. Si-Mohamed, C. Dumot, J. Kim, V. Hubert, C. C. Da Silva, M. Vandamme, E. Chereul, J. Balegamire, Y. Chevalier, Y. Berthezene, L. Boussel, P. Douek, D. P. Cormode, and M. Wiart, “Multicolor spectral photon counting ct monitors and quantifies therapeutic cells and their encapsulating scaffold in a model of brain damage,” Nanotheranostics 4, pp. 129–141, 2020.

[10] X. Chen, J. Zhang, K. Wu, X. Wu, J. Tang, S. Cui, D. Cao, R. Liu, C. Peng, L. Yu, and J. Ding, “Visualizing the in vivo evolution of an injectiable and thermosensitive hydrogel using tri-modal bioimaging,” Small 4, p. 2000310, 2020.

[11] K. M. Pawelec, T. A. Schoborg, and E. M. Shapiro, “Computed tomography technologies to measure key structural features of polymeric biomedical implants from bench to bedside,” J. Biomed. Mater. Res., pp. 1–9, 2024.

[12] T. Lex, B. R. Brummel, M. F. Attia, L. N. Giambalvo, K. G. Lee, B. A. Van Horn, D. C. Whitehead, and F. Alexis, “Iodinated polyesters with enhanced x-ray contrast properties for biomedical imaging,” Scientific Reports 10, p. 1508, 2020.

[13] K. Lei, Y. Chen, J. Wang, X. Peng, L. Yu, and J. Ding, “Non-invasive monitoring of in vivo degradation of a radiopaque termoreversible hydrogel and its efficacy in preventing post-operative adhesions,” Acta Biomaterialia 55, pp. 396–409, 2017.

14. X. Wu, X. Wang, X. Chen, X. Yang, Q. Ma, G. Xu, L. Yu, and J. Ding, “Injectable and thermosensitive hydrogels mediating a universal macromolecular contrast agent with radiopacity for noninvasive imaging of deep tissues,” Bioactive Materials 6, pp. 4717–4728, 2021.

[15] K. R. Sneha and G. S. Sailaja, “Intrinsically radiopaque biomaterial assortments: a short review on the physical principles, x-ray imageability, and state-of-the-art developments,” J. Mater. Chem. B 9, p. 8569, 2021.

[16] Y. C. Dong, M. Hajfathalian, P. S. N. Maidment, J. C. Hsu, P. C. Naha, S. Si-Mohamed, M. Breuilly, J. Kim, P. Chhour, P. Douek, H. I. Litt, and D. P. Cormode, “Effect of gold nanoparticle size on their properties as contrast agents for computed tomography,” Scientific Reports 9, p. 14912, 2019.

[17] M. Delemeester, K. M. Pawelec, J. M. L. Hix, J. R. Siegenthaler, M. Lissy, P. C. Douek, A. Houmeau, S. A. Si-Mohamed, and E. M. Shapiro, “Device design and advanced computed tomography of 3D printed radiopaque composite scaffolds and meniscus,” Advanced Functional Materials 2402404860, 2024.

[18] P. F. FitzGerald, R. E. Colborn, P. M. Edic, J. W. Lambert, A. S. Torres, P. J. Bonitatibus, and B. M. Yeh, “Ct image contrast of high-z elements: Phantom imaging studies and clinical implications,” Radiology 278(3), pp. 723–733, 2016.

[19] S. Chakravarty, J. M. L. Hix, K. A. Wiewiora, M. C. Volk, E. Kenyon, D. D. Shuboni-Mulligan, B. Blanco-Fernandez, M. Kiupel, J. Thomas, L. F. Sempere, and E. M. Shapiro, “Tantalum oxide nanoparticles as versatile contrast agents for x-ray computed tomography,” Nanoscale 12, pp. 7720–7734, 2020.

[20] J. W. Lambert, Y. Sun, C. Stillson, Z. Li, R. Kumar, S. Wang, P. F. FitzGerald, P. J. Bonitatibus, R. E. Colborn, J. C. Roberts, P. M. Edic, M. Marino, and B. M. Yeh, “An intravascular tantalum oxide-based CT contrast agent: preclinical evaluation emulating overweight and obese patient size.,” Radiology 289(1), pp. 103–110, 2018.

[21] K. Pawelec, E. Tu, S. Chakravarty, J. Hix, L. Buchanan, L. Kenney, F. Buchanan, N. Chatterjee, S. Das, A. Alessio, and E. Shapiro, “Incorporating tantalum oxide nanoparticles into implantable polymeric biomedical devices for radiological monitoring,” Adv. Healthcare Mater. 2203167, 2023.

[22] X. Li, Q. Zou, J. Wei, and W. Li, “The degradation regulation of 3d printed scaffolds for promotion of osteogenesis and in vivo tracking,” Composites Part B 222, p. 109084, 2021.

[23] E. Pamula and E. Menaszek, “In vitro and in vivo degradation of poly(l-lactide-co-glycolide) films and scaffolds,” J. Mater. Sci.: Mater. Med. 19, pp. 2063–2070, 2008.

[24] M. Bartnikowski, T. R. Dargaville, S. Ivanovski, and D. W. Hutmacher, “Degradation mechanisms of polycaprolactone in the context of chemistry, geometry and environment,” Progress in Polymer Science 96, pp. 1–20, 2019.

[25] K. M. Pawelec, J. Hix, and E. Shapiro, “Functional attachment of primary neurons and glia on radiopaque implantable biomaterials for nerve repair,” Nanomedicine: Nanotechnology, Biology, and Medicine 52, p. 102692, 2023.

[26] K. M. Pawelec, J. M. L. Hix, and E. M. Shapiro, “Material matters: Degradation products affect regenerating Schwann cells,” Biomaterials Advances 159, p. 213825, 2024.

[27] K. M. Pawelec, J. M. L. Hix, A. Troia, K. W. MacRenaris, M. Kiupel, and E. M. Shapiro, “In vivo micro-computed tomography evaluation of radiopaque, polymeric device degradation in normal and inflammatory environments,” Acta Biomaterialia 181, pp. 222–234, 2024.

[28] L. Wu and J. Ding, “In vitro degradation of three-dimensional porous poly(d,l-lactide-co-glycolide) scaffolds for tissue engineering,” Biomaterials 25, pp. 5821–5830, 2004.

[29] A. N. Smith, J. B. Ulsh, R. Gupta, M. M. Tang, A. P. Peredo, T. D. Teinturier, R. L. Mauck, S. Gullbrand, and M. W. Hast, “Characterization of degradation kinetics of additively manufactured PLGA under variable mechanical loading paradigms,” Journal of the Mechanical Beavior of Biomedical Materials 153, p. 106457, 2024.

[30] R. H. Rigdon, “Local reaction to polyurethane - a comparative study in the mouse, rat and rabbit,” J. Biomed. Mater. Res. 7, pp. 79–93, 1973.

[31] N. Artzi, N. Oliva, C. Puron, S. Shitreet, S. Artzi, A. bon Ramos, A. Groothuis, G. Sahagian, and E. R. Edelman, “In vivo and in vitro tracking of erosion in biodedgradable materials using non-invasive fluorescence imaging,” Nature Materials 10, pp. 704–709, 2011.

[32] K. Lei, Y. Chen, J. Wang, X. Peng, L. Yu, and J. Ding, “Non-invasive monitoring of in vivo degradation of a radiopaque thermoreversible hydrogel and its efficacy in preventing post-operative adhesions,” Acta Biomaterialia 55, pp. 396–409, 2017.

[33] T. R. Olsen, L. L. Davis, S. E. Nicolau, C. C. Duncan, D. C. Whitehead, B. A. Van Horn, and F. Alexis, “Non-invasive deep tissue imaging of iodine modified poly(caprolactone-co-1-4-oxepan-1,5-dione) using X-ray,” Acta Biomaterialia 20, pp. 94–103, 2015.

[34] m. Le Goas, F. Testard, O. Tache, N. Debou, B. Cambien, G. Carrot, and J.-P. Renault, “How do surface properties of nanoparticles influence their diffusion in the extracellular matrix? a model study in matrigel using polymer-grafted nanoparticles,” Langmuir 36, pp. 10460–10470, 2020.

[35] B. Reid, M. Gibson, A. Singh, J. Taube, C. Furlong, M. Murcia, and J. Elisseeff, “PEG hydrogel degradation and the role of the surrounding tissue environment,” J. Tissue Eng. Regen. Med. 9, pp. 315– 318, 2015.

[36] A. Al Shoyaib, S. R. Archie, and V. T. Karamyan, “Intraperitoneal route of drug administration: Should it be used in experimental animal studies?,” Pharm. Res. 37, p. 12, 2020.

[37] A. J. S. McCartan, D. W. Curran, and R. J. Mrsny, “Evaluating parameters affecting drug fate at the intramuscular injection site,” Journal of Controlled Release 336, pp. 322–335, 2021.

[38] E. Vey, C. Rodger, J. Booth, M. Claybourn, A. F. Miller, and A. Saiani, “Degradation kinetics of poly(lactic-co-glycolic) acid block copolymer cast films in phosphate buffer solution as revealed by infrared and Raman spectroscopies,” Polymer Degradation and Stability 96, pp. 1882–1889, 2011.

[39] N. Siddiqui, S. Asawa, B. Birru, R. Baadhe, and S. Rao, “PCL-based composite scaffold matrices for tissue engineering applications,” Molecular Biotechnology 60, pp. 506–532, 2018.

[40] G. Narayanan, V. N. Vernekar, E. L. Kuyinu, and C. T. Laurencin, “Poly(lactic acid)-based biomaterials for orthopaedic regenerative engineering,” Advanced Drug Delivery Reviews 107, pp. 247–276, 2016.

[41] J. Meganck and B. Liu, “Dosimetry in micro-computed tomography: a review of the measurement methods, impacts, and characterization of the quantum gx,” Mol Imaging Biol 19, pp. 499–511, 2017.

[42] N. Ziemkiewicz, G. Hilliard, N. A. Pullen, and K. Garg, “The role of innate and adaptive immune cells in skeletal muscle regeneration,” International Journal of Molecular Sciences 22, p. 3265, 2021.

[43] P. Habibovic and K. de Groot, “Osteoinductive biomaterials - properties and relevance in bone repair,” J. Tissue Eng. Regen. Med. 1, pp. 25–32, 2007.

[44] H. Zhu, X. Ji, H. Guan, L. Zhao, L. Zhao, C. Liu, C. Cai, W. Li, T. Tao, J. Reseland, H. Haugen, and J. Xiao, “Tantalum nanoparticles reinforced polyetheretherketone shows enhanced bone formation,” Materials Science and Engineering C 101, pp. 232–242, 2019.

[45] Y. Chen, Z. Luo, W. Meng, K. Liu, Q. Chen, Y. Cai, Z. Ding, C. Huang, Z. Zhou, M. Jiang, and L. Zhou, “Decoding the “fingerprint” of implant materials: Insights into the foreign body reaction,” Small 20, p. 10325, 2024.

[46] H.-J. Sung, C. Meredith, C. Johnson, and Z. S. Galis, “The effect of scaffold degradation rate on three dimensional cell growth and angiogenesis,” Biomaterials 25, pp. 5735–5742, 2004.

[47] C. Maduka, O. Habeeb, M. Kuhnert, M. Hakun, S. Goodman, and C. Contag, “Glycolytic reprogramming underlies immune cell activation by polyethylene wear particles,” Biomaterials Advances 152, p. 213495, 2023.

48. J. V. D. Perez, B. Singhana, J. Damasco, L. Lu, P. Behlau, R. D. Rojo, E. M. Whitley, F. Heralde, A. Melancon, S. Huang, and M. P. Melancon, “Radiopaque scaffolds based on electrospun iodixanol/ polycaprolactone fibrous composites,” Materialia 14, p. 100874, 2020.

[49] C. Yang, C. Zhao, S. Wang, X. and Mengchao, Y. Zhu, L. Jing, C. Wu, and J. Chang, “Stimulation of osteogenesis and angiogenesis by micro/nano hierarchical hydroxyapatite via magrophage immunomodulation,” Nanoscale 11, p. 17699, 2019.

[50] N. Hoshyar, S. Gray, H. Han, and G. Bao, “The effect of nanoparticle size on in vivo pharmacokinetics and cellular interaction,” Nanomedicine 11(6), pp. 673–692, 2016.

[51] P. J. Bonitatibus, A. S. Torres, B. Kandapallil, B. D. Lee, G. D. Goddard, R. E. Colborn, and M. E. Marino, “Preclinical assessment of a zwitterionic tantalum oxide nanoparticle x-ray contrast agent,” ACS Nano 6(8), pp. 6650–6658, 2012.

[52] A. B. Engin, D. Nikitovic, M. Neagu, P. Henrich-Noach, A. O. Docea, M. I. Shtilman, K. Golokhvast, A. M. Tsatsakis, “Mechanistic understanding of nanoparticles’ interactions with extracellular matrix: the cell and immune system,” Particle and Fibre Technology 14, p. 22, 2017.

[53] N. Otsu, “A threshold selection method from gray-level histograms,” IEEE Transactions on Systems, Man, and Cybernetics 9(1), pp. 62–66, 1979.

[54] P. A. Yushkevich, J. Piven, H. C. Hazlett, R. G. Smith, S. Ho, J. C. Gee, and G. Gerig, “User-guided 3D active contour segmentation of anatomical structures: Significantly improved efficiency and reliability,” NeuroImage 31, pp. 1116–1128, 2006.

