## Supplemental for "Material Composition and Implantation Site Affect in vivo Device Degradation Rate"

*Kendell M Pawelec, Jeremy ML Hix, Arianna Troia, Matti Kiupel, Erik M Shapiro**

### S1. $TaO_x$ Nanoparticle Characterization

Hydrophobic $TaO_x$ nanoparticles were manufactured based on our previous procedure with one minor change.[11] Hydrophobicity was imparted by coating nanoparticles with hexadecyltriethoxysilane (HDTES, Gelest Inc, cat no SIH5922.0). The $TaO_x$ nanoparticles were imaged via transmission electron microscopy, **Figure S1**. They were spherical particles, with a mean diameter of 6.5 ± 3nm when analyzed via Image J.

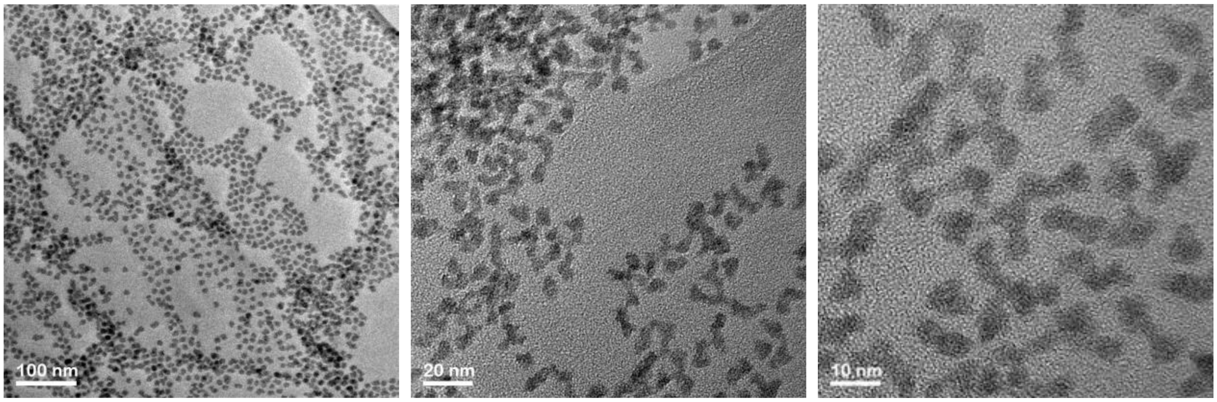


**Figure S1**. Hydrophobic $TaO_x$ nanoparticles visualized via transmission electron microscopy. At all magnifications, the particles show a tight distribution in size (3-9nm).

The mass attenuation coefficient for a phantom of polycaprolactone (PCL) with 20wt% $TaO_x$ nanoparticles was calculated using the NIST FFast Physical Reference Database (nist.gov), **Figure S2**. The attenuation coefficient was dominated by the $TaO_x$ present. No difference in the coefficient is expected with different polymer matrices, as all are composed of the same basic elements.

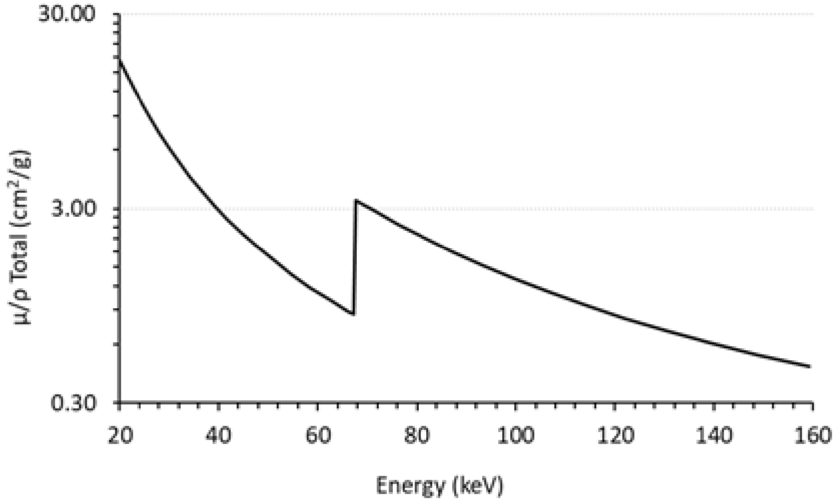

**Figure S2.** Mass attenuation coefficient calculated for polycaprolactone (PCL) with 20wt% $TaO_x$ phantom. The coefficient is dominated by the $TaO_x$ nanoparticles within the imaging range.

#### S.2 Composite Characterization

Thermogravimetric analysis (TGA) was used to characterize the weight percentage of $TaO_x$ incorporated into polymeric films and devices. Using a TA Q500 (TA Instruments), 12-14 mg of dry film or device was weighed in an alumina pan. The sample was heated from 20 - 600°C at a rate of 10°C/min under a nitrogen environment. The weight percentage of nanoparticles was calculated as the difference between the initial and final mass. A final mass of < 2wt% was considered indicative of successful removal of sucrose and salt in the washing step. Results of TGA characterization are presented in **Table S1**.

#### S3. Phantom Characterization

Pore size was analyzed from cross-sections of the phantoms, obtained via micro-computed tomography (µCT). All tomography images were obtained using a Perkin-Elmer Quantum GX. Groups of three phantoms were imaged in air, at 90 keV, 88 µA, with a 25 mm field of view at a 50 µm resolution.

To ensure the same process was applied to all percentages of $TaO_x$ incorporation, Image J was used. Cross-sections (3 per phantom) were binarized using Weka trainable Segmentation (a built-in Plug-in). The binary images were then prepared for quantification via processing tools built into Image J: "despeckle" was applied to reduce noise, followed by the application of a watershed to close interconnected pores. The size of the segmented pores was then quantified by their corresponding Feret's diameter. The mean and median of pores sizes for each group is reported in **Table S1** (n=3 scaffolds per group). No statistical differences in the mean pore size were found. Variability in the mean pore size measured was higher with lower incorporation of $TaO_x$ (≤ 5wt%), attributed to greater error in the segmentation procedure. Electron microscopy images taken of cross-sections of the porous phantoms confirmed a uniform porous morphology, **Figure S3**.

**Table S1. Phantom features with varying composition (mean ± standard deviation)**

| Polymer Matrix | Nominal $TaO_x$ (wt%) | Measured $TaO_x$ (wt%) | Mean Pore Size (µm) | X-Ray Attenutation (HU) |
|---|---|---|---|---|
| PCL | 5 | 7.12 ± *1.0* | 358 ± *58.5* | 352.0 ± *55.5* |
| | 20 | 17.61 ± *2.3* | 395 ± *11.2* | 812.9 ± *158* |
| PLGA 85:15 | 0 | 1.00 ± *0.5* | 420 ± *33.8* | 126.8 ± *48.3* |
| | 5 | 5.65 ± *1.3* | 336 ± *13.4* | 298.2 ± *62.7* |
| | 20 | 16.35 ± *3.4* | 373 ± *10.3* | 782.6 ± *118* |
| PLGA 50:50 | 5 | 5.16 ± *1.8* | 329 ± *15.7* | 375.9 ± *49.4* |
| | 20 | 16.55 ± *2.0* | 351 ± *7.98* | 814.4 ± *108* |

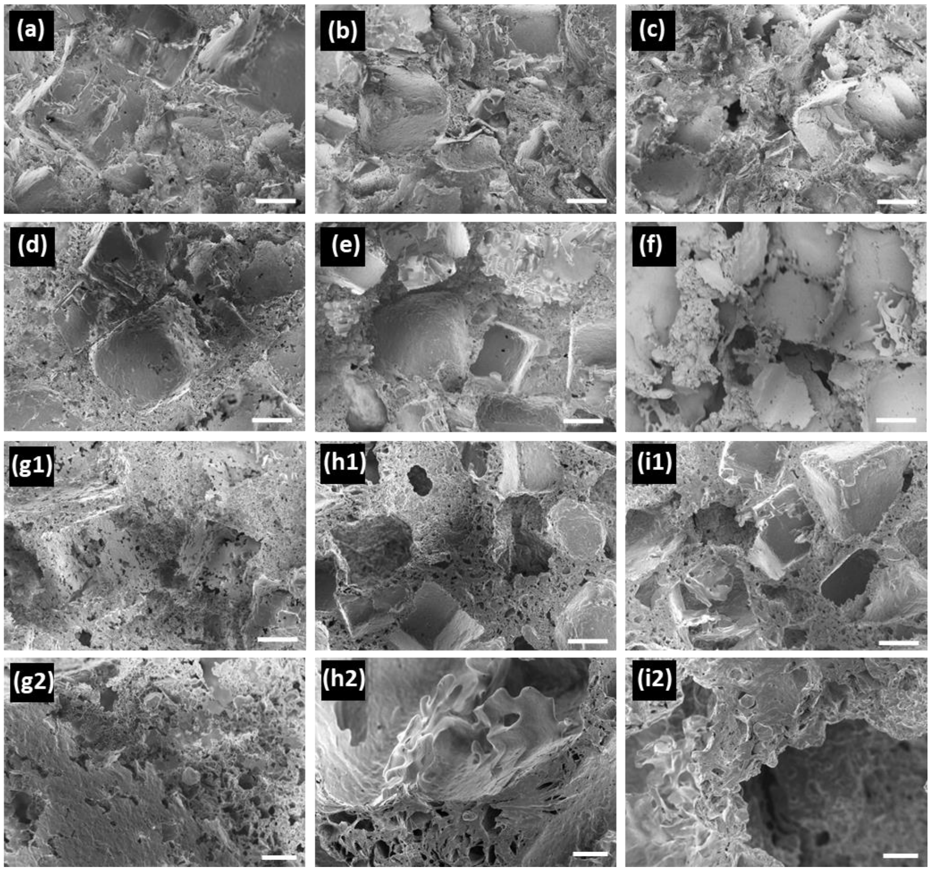


**Figure S3.** Phantom morphology did not change with addition of $TaO_x$ nanoparticles up to 20wt%, as observed in scanning electron microscopy (SEM): (a-c) PLGA 85:15, (d-f) PLGA 50:50, (g-i) PCL with (a, d, g) 0wt% $TaO_x$, (b, e, h) 5wt% $TaO_x$, and (c, f, i) 20wt% $TaO_x$. All have macropores (>200µm) with microporous walls, seen more clearly at high magnification (g2-i2). Scale bar (a-f, g1-i1): 100µm, (g2-i2): 20µm.

**S.4 Tissue Pathology**

Tissue samples of the heart, brain, kidney, liver, and spleen were also collected post-euthanasia for histopathological hematoxylin and eosin (H&E) staining to investigate for signs of adverse systemic reactions to either the radiation exposure from CT scanning or the presence of nanoparticles. No signs of pathology were detected due to the devices or nanoparticles. A representative panel of tissues is presented in **Figure S4**.

Representative histological sections around the implanted phantoms were stained with H&E at 20 weeks post-implantation. These were examined for measures of inflammation, fibrosis, and mineralization by a certified tissue pathologist. Inflammation at the site was characterized by granulomatous cellulitis. The presence of phantoms increased inflammatory reactions at both implantation sites, with the most sustained reactions occurring around PCL phantoms, **Figure S5**.

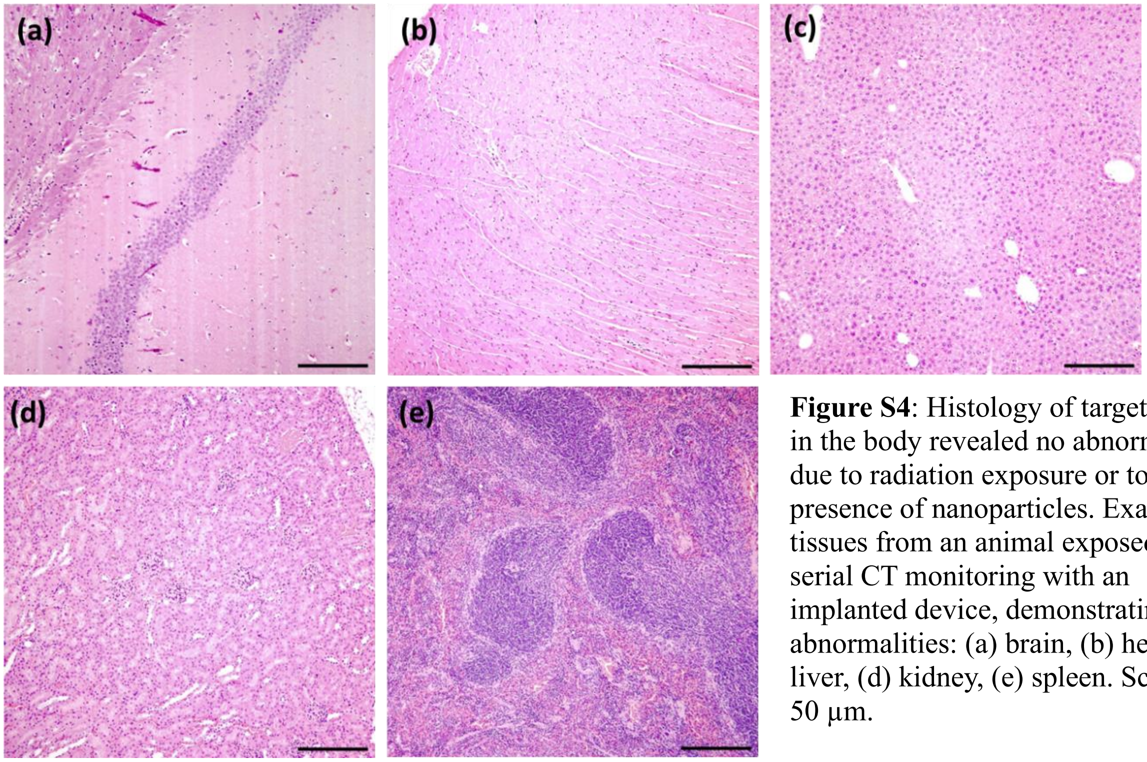


**Figure S4**: Histology of target tissues in the body revealed no abnormalities due to radiation exposure or to the presence of nanoparticles. Example tissues from an animal exposed to serial CT monitoring with an implanted device, demonstrating no abnormalities: (a) brain, (b) heart, (c) liver, (d) kidney, (e) spleen. Scale bar: 50 µm.

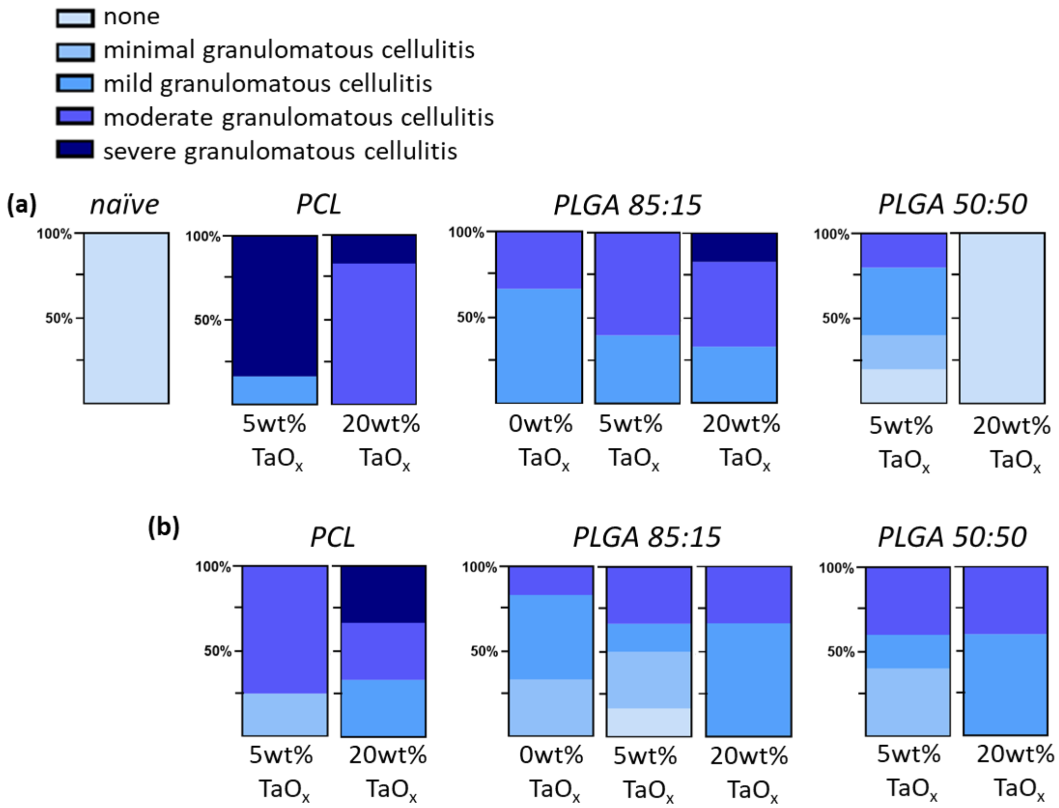


**Figure S5.** Inflammation, measured by granulomatous cellulitis, around implanted phantoms after 20 weeks. Phantoms were implanted (a) intramuscularly and (b) intraperitoneally. Each phantom was composed of a matrix polymer and the incorporation of $TaO_x$ nanoparticles up to 20wt%. Naïve tissue was included as a control.

#### S.5 In vivo study: Blood Analysis

At the end of the study, following euthanasia, blood was collected from all animals. The blood was used for complete blood cell count (CBC) and clinical pathology. In the CBC analysis, no significant changes were noted in the white blood cell count (WBC), although significant differences in relative amount of blood components, such as monocytes and lymphocytes was noted, **Table S2**. Changes in blood cell count were not linked to μCT scanning or incorporation of $TaO_x$ nanoparticles.

**Table S2** Complete blood count of all groups; naïve animals are reference.

| UNIT | HEMATOLOGY CBC Automated | Reference | PLGA 85:15 0wt% $TaO_x$ IM* | PLGA 85:15 0wt% $TaO_x$ IP* | PCL 5wt% $TaO_x$ IM | PCL 5wt% $TaO_x$ IP | PCL 20wt% $TaO_x$ IM | PCL 20wt% $TaO_x$ IP | PLGA 50:50 5wt% $TaO_x$ IM | PLGA 50:50 5wt% $TaO_x$ IP | PLGA 50:50 20wt% $TaO_x$ IM | PLGA 50:50 20wt% $TaO_x$ IP | PLGA 85:15 5wt% $TaO_x$ IM | PLGA 85:15 5wt% $TaO_x$ IP | PLGA 85:15 20wt% $TaO_x$ IM | PLGA 85:15 20wt% $TaO_x$ IP |
|---|---|---|---|---|---|---|---|---|---|---|---|---|---|---|---|---|
| **g dL$^{-1}$** | Total Protein (Refractometry) | 6.73 ± 0.5 | 6.07 ± 0.2 | 5.78 ± 0.2 | 6.05 ± 0.1 | 5.98 ± 0.2 | 6.25 ± 0.1 | 6.07 ± 0.3 | 6.5 ± 0.1 | 6.26 ± 0.1 | 6.28 ± 0.3 | 6.22 ± 0.3 | 6.10 ± 0.2 | 5.94 ± 0.2 | 6.03 ± 0.2 | 5.90 ± 0.2 |
| **10x10$^{6}$ μL$^{-1}$** | RBC | 10.40 ± 0.4 | 9.63 ± 0.2 | 9.73 ± 0.4 | 9.37 ± 0.3 | 9.63 ± 0.1 | 9.78 ± 0.2 | 9.28 ± 0.4 | 9.8 ± 0.2 | 9.46 ± 0.2 | 9.27 ± 0.3 | 9.12 ± 0.6 | 9.67 ± 0.3 | 9.70 ± 0.3 | 9.53 ± 0.2 | 9.44 ± 0.2 |
| **g dL$^{-1}$** | Hgb | 16.55 ± 0.9 | 14.50 ± 1.0 | 15.15 ± 0.4 | 15.17 ± 0.4 | 15.40 ± 0.2 | 15.92 ± 0.3 | 14.82 ± 0.8 | 15.8 ± 0.1 | 15.26 ± 0.3 | 15.13 ± 0.3 | 14.54 ± 1.0 | 15.55 ± 0.3 | 15.40 ± 0.4 | 15.35 ± 0.3 | 15.04 ± 0.4 |
| **%** | Hct | 50.25 ± 2.6 | 45.67 ± 1.1 | 46.67 ± 1.5 | 45.67 ± 1.7 | 46.00 ± 0.7 | 48.60 ± 1.6 | 46.00 ± 1.5 | 48 ± 1.0 | 46.80 ± 1.2 | 46.00 ± 0.6 | 44.60 ± 3.6 | 48.00 ± 0.6 | 47.80 ± 1.6 | 47.00 ± 1.0 | 46.40 ± 1.0 |
| **%** | HCT Spun | 48.25 ± 2.9 | 44.17 ± 1.8 | 45.33 ± 1.8 | 45.67 ± 0.9 | 46.50 ± 1.1 | 48.40 ± 1.0 | 44.80 ± 1.3 | 47 ± 1.0 | 45.60 ± 0.5 | 44.67 ± 1.1 | 42.60 ± 4.0 | 46.50 ± 1.5 | 46.80 ± 1.5 | 45.17 ± 0.7 | 45.80 ± 1.2 |
| **fL** | MCV | 48.25 ± 0.8 | 47.33 ± 0.5 | 47.83 ± 1.2 | 48.83 ± 0.9 | 48.00 ± 0.7 | 49.80 ± 0.4 | 49.67 ± 1.2 | 49 ± 1.0 | 49.60 ± 0.8 | 49.83 ± 1.6 | 48.80 ± 0.7 | 49.83 ± 0.9 | 49.60 ± 0.5 | 49.17 ± 0.4 | 49.00 ± 0.6 |
| **pg** | MCH | 16.00 ± 0.0 | 15.67 ± 0.5 | 15.83 ± 0.4 | 16.17 ± 0.4 | 16.00 ± 0.0 | 16.20 ± 0.4 | 16.00 ± 0.0 | 16 ± 0.0 | 16.20 ± 0.4 | 16.33 ± 0.5 | 16.00 ± 0.0 | 21.17 ± 12 | 16.00 ± 0.0 | 16.00 ± 0.0 | 16.00 ± 0.0 |
| **g dL$^{-1}$** | MCHC | 33.00 ± 0.5 | 32.67 ± 0.5 | 32.67 ± 0.5 | 33.33 ± 0.9 | 33.50 ± 0.5 | 32.40 ± 0.8 | 32.00 ± 0.6 | 33 ± 1.0 | 32.60 ± 0.5 | 33.00 ± 0.8 | 32.40 ± 0.5 | 32.33 ± 0.5 | 32.00 ± 0.0 | 32.67 ± 0.5 | 32.80 ± 0.4 |
| **g dL$^{-1}$** | CHCM | 30.00 ± 0.8 | 30.83 ± 0.4 | 30.50 ± 0.5 | 31.33 ± 0.9 | 31.00 ± 0.0 | 30.20 ± 0.4 | 30.17 ± 0.7 | 30.5 ± 0.5 | 29.80 ± 0.4 | 30.50 ± 0.5 | 30.40 ± 0.5 | 30.17 ± 0.4 | 30.00 ± 0.0 | 30.33 ± 0.5 | 30.20 ± 0.4 |
| **%** | RDW | 14.00 ± 0.0 | 13.17 ± 0.4 | 13.83 ± 0.7 | 14.33 ± 0.7 | 14.00 ± 0.0 | 14.00 ± 0.0 | 15.67 ± 1.8 | 14 ± 0.0 | 14.00 ± 0.0 | 14.17 ± 0.4 | 14.60 ± 0.8 | 14.17 ± 0.4 | 14.00 ± 0.0 | 14.00 ± 0.0 | 14.00 ± 0.0 |
| **%** | Total Retic Pct | 3.48 ± 0.4 | 3.53 ± 0.2 | 3.55 ± 0.4 | 4.67 ± 0.7 | 3.85 ± 0.2 | 4.06 ± 0.1 | 5.65 ± 1.4 | 3.9 ± 0.2 | 4.02 ± 0.4 | 4.00 ± 0.4 | 4.78 ± 1.2 | 3.92 ± 0.4 | 3.96 ± 0.2 | 4.07 ± 0.3 | 3.96 ± 0.3 |
| **10x10$^{4}$ μL$^{-1}$** | Total Retic # | 35.92 ± 3.1 | 34.09 ± 2.4 | 34.60 ± 4.3 | 43.49 ± 5.0 | 37.18 ± 1.5 | 39.58 ± 1.9 | 51.97 ± 10 | 38.19 ± 1.3 | 38.05 ± 3.9 | 37.10 ± 3.6 | 43.08 ± 9.4 | 42.82 ± 11 | 38.43 ± 2.8 | 38.83 ± 3.0 | 37.69 ± 3.9 |
| **pg** | CHr | 16.00 ± 0.1 | 15.93 ± 0.2 | 15.90 ± 0.3 | 16.48 ± 0.2 | 16.23 ± 0.2 | 16.44 ± 0.1 | 16.50 ± 0.5 | 16.35 ± 0.2 | 16.20 ± 0.2 | 16.68 ± 0.4 | 16.24 ± 0.3 | 16.23 ± 0.2 | 16.14 ± 0.1 | 16.12 ± 0.2 | 15.92 ± 0.2 |
| **10x10$^{3}$ μL$^{-1}$** | Platelet | 1012.8 ± 62 | 1155.5 ± 31 | 1041.8 ± 192 | 1012.8 ± 55 | 911.3 ± 82 | 1025.0 ± 51 | 1263.3 ± 180 | 772.5 ± 58 | 940.6 ± 50 | 821.7 ± 284 | 725.8 ± 344 | 898.3 ± 91 | 920.0 ± 50 | 955.3 ± 45 | 866.0 ± 108 |
| **fL** | MPV | 5.63 ± 0.5 | 5.43 ± 0.4 | 5.97 ± 0.7 | 5.33 ± 0.2 | 5.75 ± 0.7 | 5.66 ± 0.4 | 6.22 ± 1.1 | 7.5 ± 0.8 | 6.16 ± 0.7 | 6.68 ± 1.0 | 7.38 ± 2.1 | 5.18 ± 0.2 | 5.70 ± 0.3 | 5.25 ± 0.2 | 5.74 ± 0.5 |
| **%** | Pct | 0.60 ± 0.1 | 0.63 ± 0.1 | 0.60 ± 0.1 | 0.55 ± 0.1 | 0.50 ± 0.1 | 0.60 ± 0.1 | 0.78 ± 0.1 | 0.6 ± 0.0 | 0.60 ± 0.1 | 0.52 ± 0.1 | 0.48 ± 0.2 | 0.45 ± 0.1 | 0.52 ± 0.0 | 0.48 ± 0.0 | 0.52 ± 0.0 |
| **10x10$^{3}$ μL$^{-1}$** | WBC | 7.40 ± 0.9 | 5.78 ± 1.3 | 5.97 ± 0.9 | 5.32 ± 1.7 | 4.63 ± 0.6 | 4.74 ± 1.6 | 5.73 ± 1.1 | 4.85 ± 0.9 | 4.52 ± 0.5 | 4.93 ± 2.1 | 4.54 ± 1.4 | 6.18 ± 0.9 | 5.20 ± 1.2 | 5.93 ± 1.9 | 4.74 ± 1.0 |
| **MANUAL** | **HEMATOLOGY CBC Manual Differential Counts** | | | | | | | | | | | | | | | |
| **10x10$^{3}$ μL$^{-1}$** | Seg Neut | 1.33 ± 0.2 | 1.30 ± 0.5 | 1.75 ± 0.6 | 0.90 ± 0.3 | 1.18 ± 0.2 | 0.88 ± 0.3 | 1.78 ± 0.7 | 1.15 ± 0.1 | 1.30 ± 0.2 | 1.12 ± 0.3 | 1.46 ± 0.6 | 0.88 ± 0.2 | 1.12 ± 0.3 | 1.35 ± 0.5 | 1.34 ± 0.3 |
| **10x10$^{3}$ μL$^{-1}$** | Band Neut | 0.00 ± 0.0 | 0.00 ± 0.0 | 0.00 ± 0.0 | 0.00 ± 0.0 | 0.00 ± 0.0 | 0.00 ± 0.0 | 0.00 ± 0.0 | 0 ± 0.0 | 0.00 ± 0.0 | 0.00 ± 0.0 | 0.00 ± 0.0 | 0.00 ± 0.0 | 0.00 ± 0.0 | 0.00 ± 0.0 | 0.00 ± 0.0 |
| **10x10$^{3}$ μL$^{-1}$** | Lymphocyte | 5.88 ± 0.6 | 4.25 ± 0.9 | 3.90 ± 0.8 | 4.12 ± 1.2 | 3.23 ± 0.4 | 3.44 ± 1.2 | 3.42 ± 0.8 | 3.25 ± 0.7 | 2.94 ± 0.4 | 3.35 ± 1.8 | 2.84 ± 0.9 | 4.95 ± 0.8 | 3.76 ± 1.1 | 4.10 ± 1.3 | 3.26 ± 0.9 |
| **10x10$^{3}$ μL$^{-1}$** | Monocyte | 0.13 ± 0.1 | 0.12 ± 0.1 | 0.28 ± 0.1 | 0.22 ± 0.2 | 0.13 ± 0.1 | 0.24 ± 0.1 | 0.37 ± 0.2 | 0.3 ± 0.2 | 0.24 ± 0.1 | 0.43 ± 0.2 | 0.20 ± 0.1 | 0.30 ± 0.1 | 0.32 ± 0.1 | 0.35 ± 0.1 | 0.06 ± 0.1 |
| **10x10$^{3}$ μL$^{-1}$** | Eosinophil | 0.10 ± 0.1 | 0.12 ± 0.1 | 0.07 ± 0.1 | 0.08 ± 0.1 | 0.08 ± 0.0 | 0.16 ± 0.1 | 0.13 ± 0.1 | 0.15 ± 0.1 | 0.04 ± 0.1 | 0.08 ± 0.0 | 0.06 ± 0.1 | 0.03 ± 0.1 | 0.06 ± 0.1 | 0.12 ± 0.1 | 0.08 ± 0.1 |
| **10x10$^{3}$ μL$^{-1}$** | Basophil | 0.00 ± 0.0 | 0.00 ± 0.0 | 0.00 ± 0.0 | 0.00 ± 0.0 | 0.00 ± 0.0 | 0.00 ± 0.0 | 0.00 ± 0.0 | 0 ± 0.0 | 0.00 ± 0.0 | 0.00 ± 0.0 | 0.00 ± 0.0 | 0.00 ± 0.0 | 0.00 ± 0.0 | 0.00 ± 0.0 | 0.00 ± 0.0 |
| **%** | Seg Neut Pct Manual | 18.25 ± 1.6 | 22.17 ± 4.6 | 29.33 ± 8.6 | 17.00 ± 5.1 | 25.25 ± 2.5 | 18.40 ± 1.4 | 31.00 ± 9.8 | 24 ± 5.0 | 28.80 ± 4.6 | 26.50 ± 11 | 30.40 ± 9.9 | 14.17 ± 2.9 | 21.60 ± 3.5 | 22.83 ± 3.5 | 29.00 ± 5.0 |
| **%** | Band Neut Pct Manual | 0.00 ± 0.0 | 0.00 ± 0.0 | 0.00 ± 0.0 | 0.00 ± 0.0 | 0.00 ± 0.0 | 0.00 ± 0.0 | 0.00 ± 0.0 | 0 ± 0.0 | 0.00 ± 0.0 | 0.00 ± 0.0 | 0.00 ± 0.0 | 0.00 ± 0.0 | 0.00 ± 0.0 | 0.00 ± 0.0 | 0.00 ± 0.0 |
| **%** | Lymphocyte Pct Manual | 78.75 ± 1.3 | 74.17 ± 4.0 | 65.17 ± 8.9 | 77.83 ± 4.7 | 70.00 ± 3.4 | 72.80 ± 3.1 | 60.17 ± 10 | 67.5 ± 1.5 | 64.80 ± 2.8 | 62.00 ± 15 | 63.80 ± 8.7 | 80.33 ± 3.1 | 71.40 ± 5.1 | 69.17 ± 4.2 | 68.00 ± 5.5 |
| **%** | Monocyte Pct Manual | 1.75 ± 1.7 | 1.67 ± 1.5 | 4.67 ± 1.2 | 3.67 ± 2.1 | 3.25 ± 1.5 | 5.40 ± 1.4 | 6.33 ± 1.8 | 5.5 ± 2.5 | 5.00 ± 2.7 | 9.83 ± 4.0 | 4.60 ± 2.3 | 4.83 ± 1.3 | 6.00 ± 1.9 | 5.83 ± 0.9 | 1.40 ± 1.4 |
| **%** | Eosinophil Pct Manual | 1.25 ± 0.9 | 2.00 ± 1.2 | 0.83 ± 0.7 | 1.50 ± 0.5 | 1.50 ± 0.9 | 3.40 ± 2.8 | 2.50 ± 1.1 | 3 ± 1.0 | 1.40 ± 1.0 | 1.67 ± 1.4 | 1.20 ± 0.7 | 0.67 ± 0.9 | 1.00 ± 0.9 | 2.17 ± 1.8 | 1.60 ± 1.5 |
| **%** | Basophil Pct Manual | 0.00 ± 0.0 | 0.00 ± 0.0 | 0.00 ± 0.0 | 0.00 ± 0.0 | 0.00 ± 0.0 | 0.00 ± 0.0 | 0.00 ± 0.0 | 0 ± 0.0 | 0.00 ± 0.0 | 0.00 ± 0.0 | 0.00 ± 0.0 | 0.00 ± 0.0 | 0.00 ± 0.0 | 0.00 ± 0.0 | 0.00 ± 0.0 |

**Denotes that no micro-computed tomography scans were performed during the study*
